# Antagonism between blue and red light-signalling controls thallus flatness in *Marchantia polymorpha*

**DOI:** 10.1101/2025.11.10.687525

**Authors:** Johannes Roetzer, Beate Asper, Zohar Meir, Natalie Edelbacher, Zsuzsanna Mérai, Sourav Datta, Liam Dolan

## Abstract

The growth orientation of the *Marchantia polymorpha* thallus – a system of dorsiventralized, indeterminate axes – is modulated by light. We show that red and blue light act antagonistically to control thallus growth tropisms, with red light signalling promoting epinasty and blue light signalling promoting hyponasty. We found that loss-of-function mutations in the blue light receptor Mp*PHOT* led to epinasty, while loss-of-function mutations in the red light receptor Mp*PHY* resulted in hyponasty. We hypothesize that these antagonistic activities of blue and red light signalling are balanced in white light, resulting in the development of flat thalli. Using time-resolved transcriptomics, we identified genes that were rapidly induced upon light exposure. Among these genes were all six members of the *M. polymorpha BBX* gene family. Mutants harbouring loss-of-function mutations in two of the six MpBBX transcription factors developed defective thallus tropisms. Mp*bbx1* loss-of-function mutants formed hyponastic thalli, while Mp*bbx5* loss-of-function mutants developed epinastic thalli. Double mutants Mpbbx1 Mp*bbx5* grew flat, supporting the hypothesis that they function antagonistically. Together, these data indicate that phototropin-mediated blue light and phytochrome-mediated red light signalling antagonistically modulate thallus tropism, and that BBX transcription factors also act antagonistically to regulate thallus flatness.

## Introduction

External environmental stimuli modulate the orientation and growth of plant organs. Tropisms are directional responses that depend on the position of the plant in relation to the stimulus. Tropic responses include phototropism, gravitropism, hydrotropism, and thigmotropism, which respectively modulate directional growth and development in response to light, gravity, water, and mechanical stimuli. In each case, the external stimulus is sensed by a receptor, and this sensing mechanism activates a response that results in the growth changes characteristic of the tropic response. Blue and red-light receptors sense light in phototropic responses. Blue light is primarily perceived by PHOTOTROPINS (PHOT), CRYPTOCHROMES (CRY), and FLAVIN-BINDING KELCH REPEAT F-BOX PROTEINS (FKF), while red and far-red light signals are detected by PHYTOCHROMES. These photoreceptors modulate tropic growth responses in angiosperms (Goyal et al., 2013, Christie, 2007, Paik & Huq, 2019).

Leaves are determinate structures that develop near the meristems of vascular plants and are flattened with distinct adaxial and abaxial surfaces in most angiosperms. Phototropism is required for leaf flatness, and it is hypothesized that it depends on the balance between blue and red light signalling (Kozuka et al., 2013, Inoue et al., 2008, Legris, 2023). It is proposed that the activities of photoreceptors influence auxin signalling, which mediates differential cell expansion during leaf flattening (Legris et al., 2021, de Carbonnel et al., 2010, Lee et al., 2024). However, the transcriptional regulators acting downstream of the photoreceptors to control leaf flatness remain to be identified.

Analogous flat photosynthetic organs include the thallus axes of thalloid liverworts such as *Marchantia polymorpha*. The *M. polymorpha* thallus axis comprises a flattened lamina with morphologically different dorsal and ventral sides. The bifurcating thallus is indeterminate, while the leaf of vascular plants is a determinate structure that develops from the flanks of meristems. Despite the morphological similarities between leaves and thalli, the developmental regulation of flatness remains poorly understood in thalli.

Here, we describe elements of the regulatory mechanisms underlying thallus flatness in *Marchantia polymorpha*. We report that red and blue light signalling pathways act antagonistically to modulate thallus tropism and shape. We show that the red-light receptor MpPHY and the blue light receptor MpPHOT antagonistically regulate thallus flatness. We also demonstrate that two B-box transcription factors, MpBBX1 and MpBBX5, are components of the signalling network that controls thallus flatness. Our findings provide a conceptual framework linking light perception to morphogenetic output in *Marchantia polymorpha*.

## RESULTS

### Red and blue light induce opposite tropic responses in thalli (Figure 1)

Light promotes the growth of the *M. polymorpha* thallus from gemma meristems (Figure 1A). Wild type plants (Tak-1 and Tak-2 accessions) grew to approximately 155 mm^2^ (surface area) in white light, while dark-grown plants hardly grew at all, with a surface area of approximately 0.35 mm^2^ at the corresponding timepoint (Figure 1B). These data indicate that light is critical for thallus growth.

**Figure 1.**
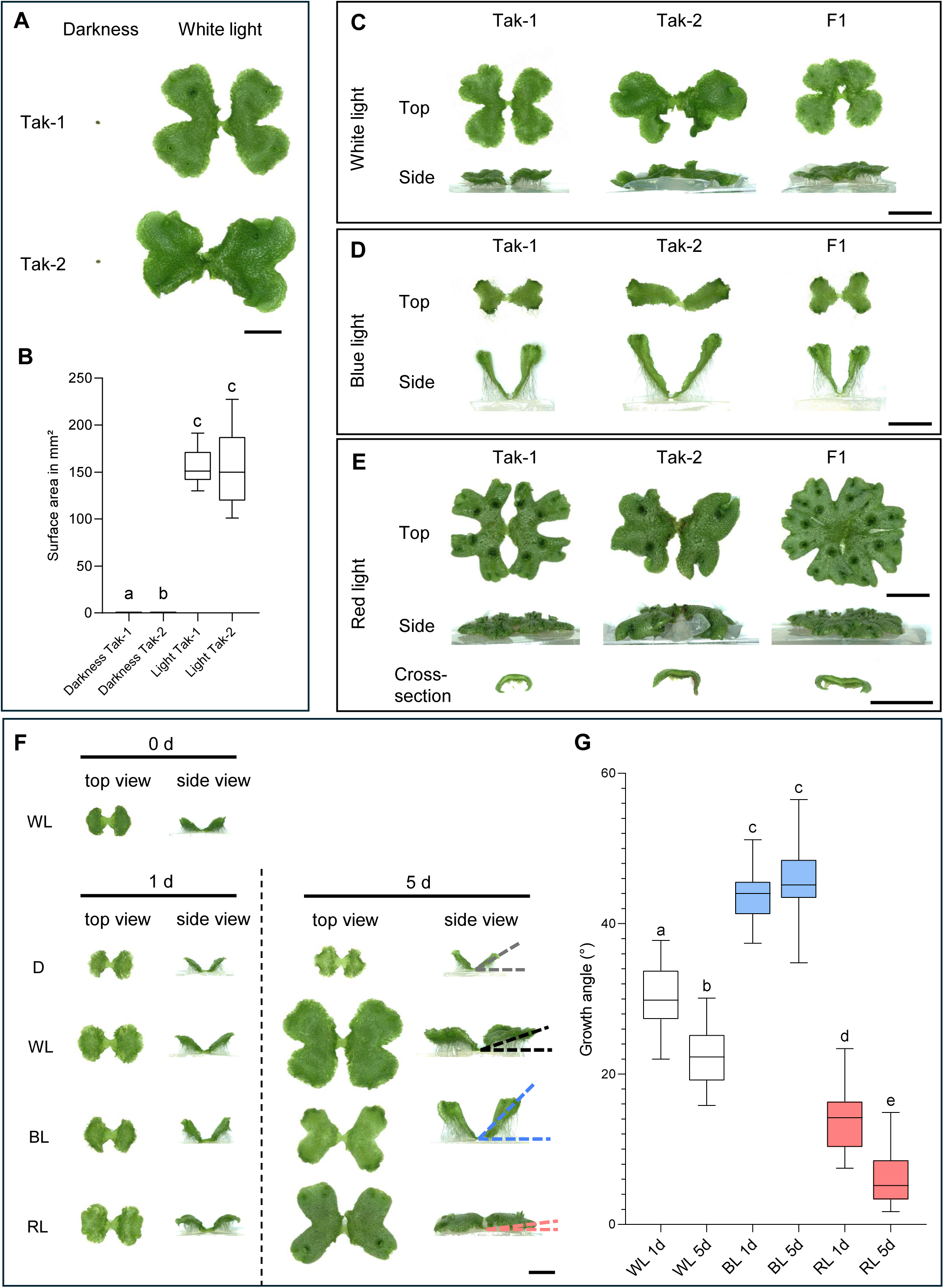
Red and blue light antagonistically control thallus tropism. (A) Light is essential for growth. Plants were grown for 14 days in continuous white light (PFD = 50-60 µmol m⁻² s⁻¹). Scale bar = 5 mm. (B) Quantification of light-vs. dark-grown plants shows that the surface area of light-grown plants was 440-fold greater than that of dark-grown plants (n = 95). Data were tested for normality using the Shapiro-Wilk test and for homogeneity of variances using Brown-Forsythe test. A Welch’s ANOVA was performed, followed by Dunnett’s T3 post-hoc test. Groups with different letters are significantly different (p < 0.05), while identical letters indicate no significant difference. (C) Wild types (Tak-1, Tak-2, and “F1,” the offspring of a cross between Tak-1 and Tak-2) grew flat in white light (14 days continuous; PFD = 50-60 µmol m⁻² s⁻¹). (D) Wild types grew hyponastically in blue light (14 days continuous; PFD = 30-40 µmol m⁻² s⁻¹). (E) Wild types grew epinastically in red light (14 days continuous; PFD = 70-80 µmol m⁻² s⁻¹). Scale bar = 1 cm for panels C–E. (F) Wavelength-specific responses after one and five days of light treatment. Scale bar = 3 mm. (G) Quantitative assay of early responses shows that measurements of the angle between the thallus tip and the agar surface revealed blue light induces hyponasty, while red light induced epinasty. Plants (n = 144) were grown for 8 days in continuous white light, incubated in darkness for one day, and then exposed to the respective light conditions for one or five days. Growth conditions were identical to those in panels C-E. Data were tested for normality using the Shapiro-Wilk test and for homogeneity of variances using Brown-Forsythe test. A one-way ANOVA was performed, followed by Tukey’s HSD post-hoc test. Groups with different letters are significantly different (p < 0.05), whereas groups with identical letters are not significantly different.

To test the effect of specific wavelengths on thallus growth, we incubated gemmae in white, blue, and red light for 14 days and compared thallus morphology (Figure 1C-E; Suppl. Figure 1E). White light-grown plants developed flat thalli growing horizontally over the growth substrate (Figure 1C; Suppl. Figure 1A-B). Blue light-grown plants developed hyponasty, where the thallus grew upward towards the light source (Figure 1D; Suppl. Figure 1E). By contrast, red light-grown plants developed epinasty, where the edges of the thallus curved downward into the growth substrate away from the light source (Figure 1E; Suppl. Figure 1E). The tropisms occurred over a range of light intensities, from approximately 1-2 µmol/m²/s to 180 µmol/m²/s, for white, blue, and red light. However, at the lowest tested light flux, plants in all wavelengths grew very slowly and remained underdeveloped (Suppl. Figure 1C). These data indicate that white, red, and blue light promote thallus growth, and that red light and blue light promote epinastic and hyponastic thallus development, respectively. This suggests that red and blue light antagonistically control the direction of thallus growth.

To further determine the effect of specific wavelengths on thallus tropism and to test if wavelength-specific responses can be activated in plants transferred from one light quality to another, we moved plants from white light into different light regimes. Gemmae were plated and grown for 8 days in white light before being transferred to darkness for 24 hours. Plants were subsequently transferred to different light conditions - darkness, full white light, blue light, or red light for a further five days. Tropic responses were measured 1 and 5 days after transfer. Thallus tropic responses were detectable as early as 1 day after the transfer from dark to light (Figure 1F). To measure thallus tropic responses, we measured the angle between the thallus tip and the agar surface, which reflected the orientation of growth at days 1 and 5 (Figure 1F-G). This angle was large when plants grew hyponastically and small when plants grew epinastically. While plants in white light grew largely horizontally, the angle decreased over time (Suppl. Figure 1D). By contrast, the angle increased in darkness, indicating that the thalli grew upward, consistent with an etiolation response (Figure 1F, Suppl. Figure 1D). The angle was greater in blue light than in all other conditions and increased between 1 and 5 days (hyponasty) (Figure 1F-G). The angle was lower in red light than in all other conditions and decreased between 1 and 5 days (epinasty) (Figure 1F-G). Furthermore, the epinasty in red light was accompanied by a downward curling of the thallus margins (Figure 1E). These data indicate that thallus tissue is responsive to changes in light regime, being hyponastic in blue light and epinastic in red light. This is consistent with our hypothesis that red and blue light act antagonistically in the control of thallus tropism.

### PHOTOTROPIN is required for blue light-mediated thallus tropism (Figure 2)

To test the hypothesis that *M. polymorpha PHOTOTROPIN* (Mp*PHOT*)is required for blue light-mediated thallus tropism and hyponasty, we generated complete loss-of-function mutants in the Mp*PHOT* gene by CRISPR/Cas9 technology using guide RNAs targeting regions near the 5’ end of the coding sequence and the start of the LOV2 domain (Suppl. Figure 2A). Protein extracts were prepared, separated on SDS-PAGE, blotted, and reacted with an MpPHOT-specific antibody. A band at approximately 124 kD was detected in wild type plants (Figure 2A) but was absent in the tested Mp*phot* mutant (Figure 2A). Three independent Mp*phot* mutant alleles, Mp*phot-3,* Mp*phot-4* and Mp*phot-5*, were used for experiments.

**Figure 2.**
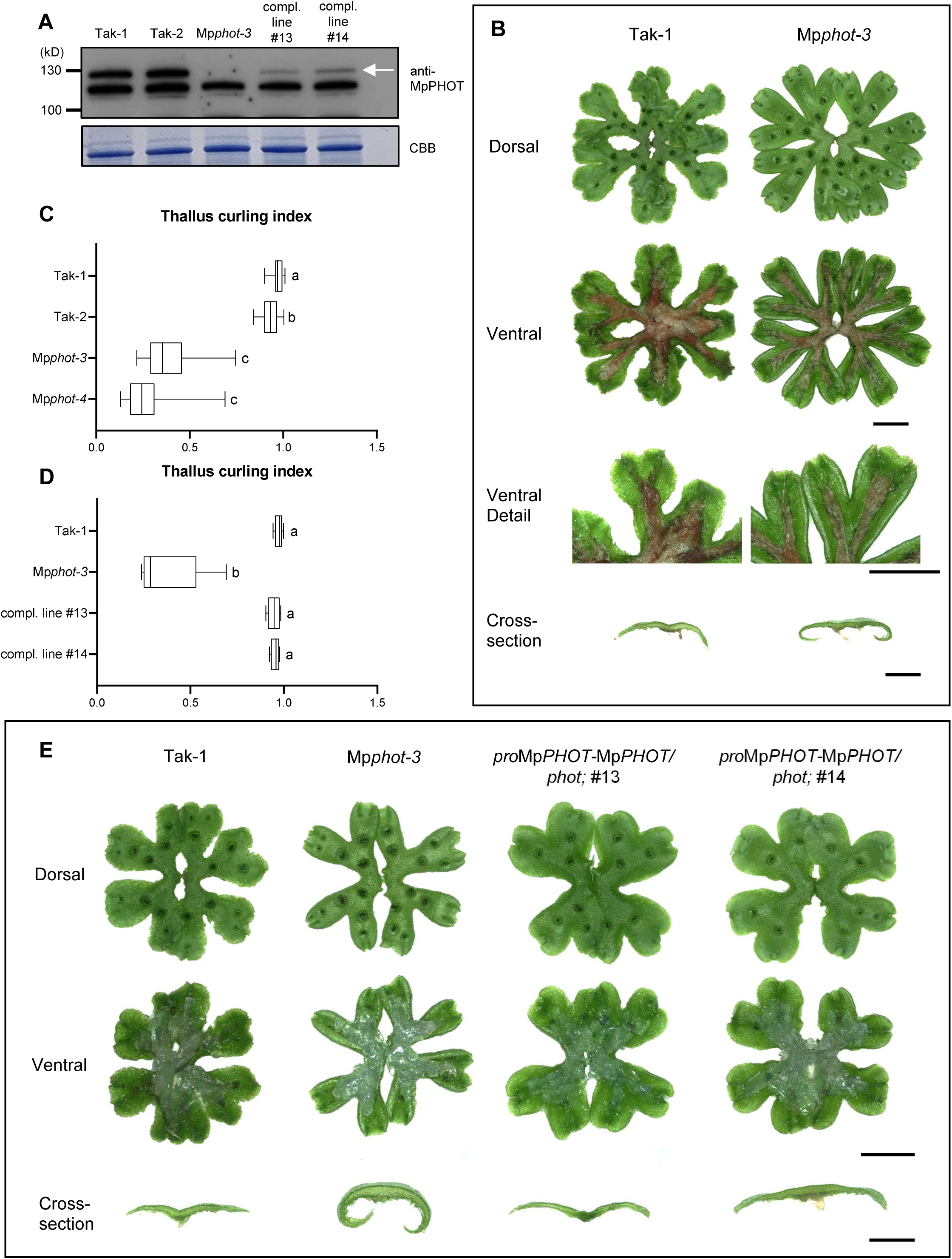
Tropic responses are defective in *phototropin* mutants. (A) MpPHOT protein was not detectable in the Mp*phot* mutant line but was detectable in the complemented Mp*phot* lines. The white arrow indicates protein accumulation at the expected size (124 kD). (B) Mp*phot* mutant lines showed defects in thallus flattening and grew epinastically. Plants were grown for 3.5 weeks under standard white light conditions. Scale bars = 10 mm for dorsal and ventral views, 2.5 mm for cross-sections. (C) Thallus curling index for cross-sections of wild type and Mp*phot* mutant lines (n = 80). Data were tested for normality using the Shapiro-Wilk test and for homogeneity of variances using Brown-Forsythe test. A Welch’s ANOVA was performed, followed by Dunnett’s T3 post-hoc test. Groups with different letters are significantly different (p < 0.05), while identical letters indicate no significant difference. (D) Thallus curling index for cross-sections of wild type, the Mp*phot* mutant line used for transformation, and complemented lines (n = 24). Data were tested for normality using the Shapiro-Wilk test and for homogeneity of variances using Brown-Forsythe test. A Welch’s ANOVA was performed, followed by Dunnett’s T3 post-hoc test. Groups with different letters are significantly different (p < 0.05), while identical letters indicate no significant difference. (E) Complemented Mp*phot* lines developed normal thallus tropism. Plants were grown for three weeks under standard white light conditions. Scale bars = 10 mm for dorsal and ventral views, 2.5 mm for cross-sections.

If Mp*PHOT* is required for hyponastic development, we predicted that Mp*phot* mutant plants would not be hyponastic. Indeed, Mp*phot* mutants under white light developed severe epinasty (Figure 2B). This suggested that in wild type, MpPHOT-mediated blue light signalling antagonizes red light-induced epinasty. In the absence of this antagonism – in Mp*phot* mutants – thalli grew in white light as though they were in monochromatic red light. We quantified these observations on epinastic growth by developing a thallus curling index. The thallus curling index is calculated as the ratio between the shortest distance between the middle of the thallus and the thallus tip and the within-tissue distance between the middle of the thallus and the thallus tip (Suppl. Figure 2B). The flatter the thallus, the closer the ratio is to 1. The average wild type thallus curling index is close to 1 (0.95), and the Mp*phot* mutant thallus curling index is 0.34 (average calculated from thallus curling index measurements in Figure 2C). The curling index of the Mp*phot* mutant in white light is similar to the curling index of wild type grown in red light (0.48) (Figure 1E; Suppl. Figure 2C). These phenotypes indicate that in the absence of Mp*PHOT* activity, thallus tropisms resemble wild type plants grown in red light, where there is no input from blue light.

To further verify that the epinasty in Mp*phot* mutants was due to the defective Mp*PHOT* gene, we complemented the Mp*phot-3* mutant with the wild-type copy of the Mp*PHOT* gene (*pro*Mp*PHOT*:Mp*PHOT*). A 124 kD protein band identified by the MpPHOT antibody accumulated in these lines (Figure 2A). The transformants developed wild-type-like thalli (Figure 2D and 2E). These data indicate that the wild type *PHOT* gene complements the defective thallus tropism phenotypes of the *phot* mutant. We thus conclude that the Mp*PHOT* gene is required for normal thallus tropism. Furthermore, the development of severe epinasty in Mp*phot* mutants grown in white light indicates that Mp*PHOT*-mediated blue light signalling acts antagonistically with red light. This suggests that a balance of red light-and blue light-signalling regulates tropic thallus growth and therefore the flatness of the *M. polymorpha* thallus.

### MpPHOT substitutes for AtPHOT1 and AtPHOT2 in blue light-mediated leaf flattening in A. thaliana (Figure 3)

PHOTOTROPIN-mediated blue light signalling promotes flattening of *A. thaliana* leaves. Leaves of At*phot1* At*phot2* double mutants are narrower and more epinastic than in wild type (Figure 3A; Suppl. Figure 3A, C) (Sullivan et al., 2008). To test if MpPHOT could substitute for At*PHOT1* or At*PHOT2* function, various Mp*PHOT* gene constructs were introduced into the At*phot1* At*phot2* double mutants by floral dip transformation. Homozygous transgenic lines were used for all complementation experiments. At*phot1* At*phot2* double mutants transformed with the Mp*PHOT* gene under the control of the At*PHOT1* promoter developed wild type phenotypes (Figure 3A; Suppl. Figure 3A, C). Similarly, At*phot1* At*phot2* double mutants transformed with the Mp*PHOT* gene under the control of At*PHOT2* promoter developed wild type phenotypes (Figure 3A; Supplementary Figure 3A, C). These observations were supported by the measured leaf flattening indices (Figure 3C). Wild-type control lines and complemented lines had leaf flattening indices of around 0.8 or higher, while in the At*phot1* At*phot2* double mutant and the negative control line (*pro*At*UBIQUITIN10*:Mp*FKF /* At*phot1* At*phot2*) it was approximately 0.4 (Figure 3C). These data indicate that expression of Mp*PHOT* from either the At*PHOT1* or At*PHOT2* promoters restored blue light-mediated leaf lamina flattening in At*phot1* At*phot2* double mutants. We independently transformed a yellow fluorescent fusion of Mp*PHOT-VENUS* into the At*phot1* At*phot2* double mutant (Figure 3A, C; Suppl. Figure 3A, C). The MpPHOT-VENUS localized to the plasma membrane in transformed *A. thaliana* plants (Figure 3E). This indicates that the MpPHOT-VENUS protein complements the At*phot1* At*phot2* double mutant phenotype and localizes to the cell surface, as it does in *M. polymorpha* (Figure 3E), consistent with previous reports (Komatsu et al., 2014).

**Figure 3.**
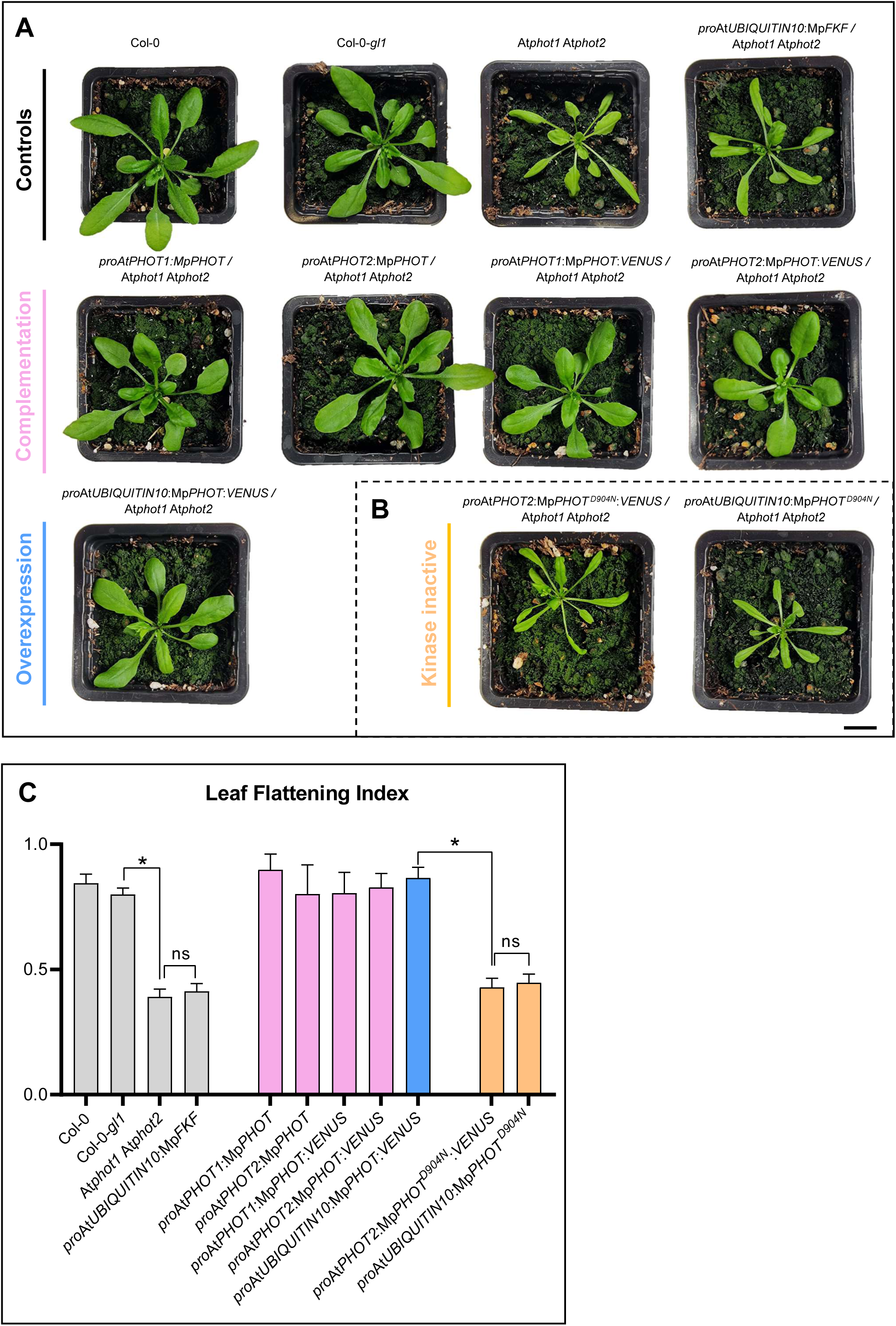
Cross-species complementation experiments reveal that *MpPHOT* function in tropism is conserved. (A) *MpPHOT* gene constructs were introduced into At*phot1* At*phot2* double mutants. Leaves of At*phot1* At*phot2* mutants were narrower and more epinastic than wild type. Expression of Mp*PHOT* from either the At*PHOT1* or At*PHOT2* promoters restored blue light-mediated leaf lamina flattening in At*phot1* At*phot2* mutants, resulting in wild type-like leaves. Col-0 and Col-0-*gl1* mutants were used as wild type controls, as the original At*phot1* At*phot2* double mutants were generated in the Col-0-*gl1* background. Independent complementation using Mp*PHOT* under the control of the At*UBIQUITIN10* promoter further confirmed that Mp*PHOT* could substitute for At*PHOT* function in leaf flattening. (B) Complementation experiments demonstrated that the kinase activity of Mp*PHOT* is essential for its function. The conserved aspartic acid residue at position 904 is required for Mp*PHOT*-mediated tropic responses. (A, B) Images show 25-day-old plants. Scale bar = 10 mm. (C) Leaf Flattening Index (LFI), calculated by dividing the surface area of the unflattened leaf by the surface area of the same leaf manually flattened. LFI was measured for leaves L3–L5 in four biological replicates per genotype, resulting in 12 LFI values per genotype (264 total measurements, 132 LFI values across 11 genotypes). Shown are the mean LFI values per genotype (x-axis) with standard deviation. A perfectly flat leaf would have an LFI of 1. Data were tested for normality using the Shapiro-Wilk test and for homogeneity of variances using Brown-Forsythe test. A Welch’s ANOVA was performed, followed by Dunnett’s T3 post-hoc test. Asterisks (*) in the comparisons denote statistically significant differences between groups (p < 0.05), while ‘ns’ indicates no significant difference (p > 0.05).

To independently test if MpPHOT could substitute for AtPHOT function, we introduced the Mp*PHOT* gene under the control of the At*UBIQUITIN10* promoter into At*phot1* At*phot2* double mutants. At*phot1* At*phot2* double mutants transformed with At*UBQ10*:Mp*PHOT* developed a flat leaf lamina indistinguishable from wild type, while At*phot1* At*phot2* double mutant leaves were epinastic (Figure 3A; Suppl. Figure 3A, C). Together, these data indicate that MpPHOT can substitute for AtPHOT1 and AtPHOT2 in *A. thaliana*.

### The conserved aspartate residue at position 904 of MpPHOT is required for organ flattening

To test if the kinase function of MpPHOT is necessary for MpPHOT-mediated signalling and development, we developed a kinase-inactive version of MpPHOT.

The aspartic acid (D) in the YRD motif at position 902-904 of the MpPHOT protein is conserved among eukaryotic kinases and required for kinase activity (Kornev et al., 2006). To generate an MpPHOT that lacks kinase activity, we mutated the aspartic acid/aspartate codon of the conserved YRD motif to YRN to an asparagine codon (D904N). This Mp*phot ^D904N^* mutant gene was placed under the control of the AtPHOT2 promoter and was introduced into the At*phot1* At*phot2* double mutant.

While plants complemented with the At*PHOT1*:Mp*PHOT* or At*PHOT2*:Mp*PHOT* constructs developed a flat leaf lamina like wild type, At*phot1* At*phot2* double mutants transformed with the Mp*phot^D904N^* mutant gene developed hyponastic leaves characteristic of the At*phot1* At*phot2* double mutants (Figure 3A, B, C; Suppl. Figure 3B, D). These data indicate that the Mp*phot ^D904N^* mutant does not complement the At*phot1* At*phot2* double mutant phenotype and that MpPHOT kinase function is required to restore the wild type leaf phenotype.

To independently verify this result, we generated At*phot1* At*phot2* double mutants transformed with wild type Mp*PHOT* and Mp*phot ^D904N^* mutant transgenes under the control of the At*UBIQUITIN10* promoter. At*phot1* At*phot2* double mutants transformed with the wild-type gene developed flat leaves and double mutants transformed with the Mp*phot ^D904N^* mutant gene developed epinastic leaves characteristic of the At*phot1* At*phot2* double mutant (Figure 3A-C; Suppl. Figure 3B, D). These data indicate that the conserved aspartate residue at position 904 is required for MpPHOT function. Leaf flattening indices of the lines transformed with the kinase-inactive MpPHOT version were similar to the At*phot1* At*phot2* double mutant and the negative control line (Fig 3C).

### PHYTOCHROME represses hyponastic thallus tropism (Figure 4)

We hypothesized that blue and red light act antagonistically in thallus tropisms. If red and blue light act antagonistically, we would predict that mutants defective in red light signalling would develop a phenotype like wild type plants grown in blue light – extreme hyponasty. To test this hypothesis, we characterized the thallus phenotypes of Mp*phy* mutants (Streubel et al., 2023). Two independent Mp*phy* mutants developed hyponastic thalli, while wild types were flat (Figure 4A). The growth angles of Mp*phy* mutants were significantly larger than wild type (Figure 4C).

**Figure 4.**
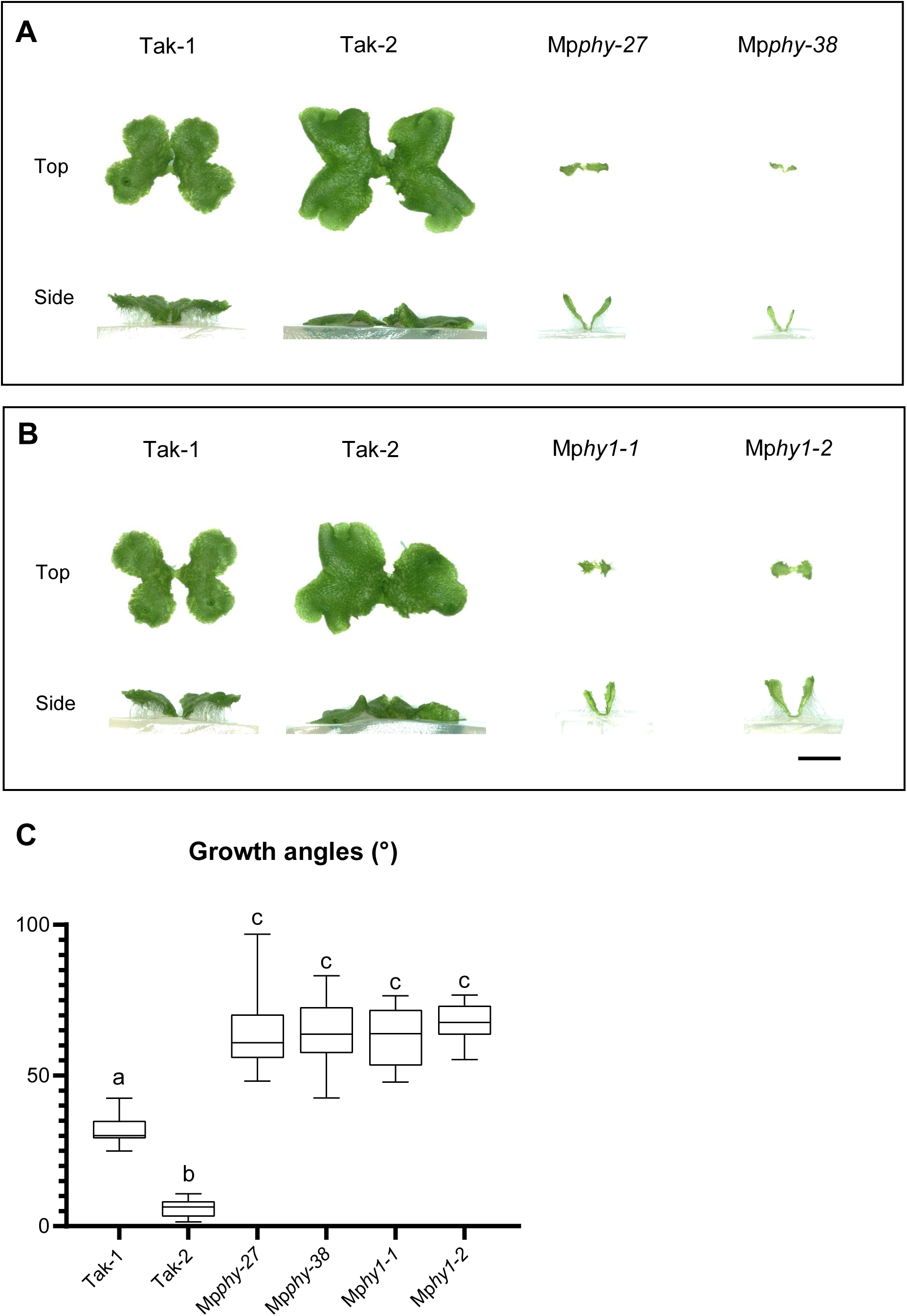
Tropic responses are defective in mutants lacking phytochrome function. (A) Mp*phy* mutants displayed strong hyponastic thallus growth, while wild type plants grew flat under standard white light conditions. (B) Mp*hy1* mutants, defective in chromophore biosynthesis, also showed hyponastic thalli. (A, B) Scale bar = 5 mm. (C) Quantification of growth angles in Mp*phy* and Mp*hy1* mutants confirmed strong hyponasty compared to wild type (n = 92). These results supported that phytochrome-mediated red light signaling inhibits hyponastic growth. Data were tested for normality using the Shapiro-Wilk test and for homogeneity of variances using Brown-Forsythe test. A Welch’s ANOVA was performed, followed by Dunnett’s T3 post-hoc test. Groups with different letters are significantly different (p < 0.05), while identical letters indicate no significant difference.

Consistent with the hyponastic phenotype of Mp*phy* loss-of-function mutants, ectopic overexpression of phytochrome (*pro*Mp*EF1*α:Mp*PHY*) caused severe epinasty; the thallus curling index of *pro*Mp*EF1*α:Mp*PHY* lines was 0.41, while the thallus curling index of wild type was close to 0.94 (Suppl. Figure 4A and B).

To independently verify that defective phytochrome signalling causes thallus hyponasty, we generated two independent alleles in the *ELONGATED HYPOCOTYL1* (*HY1*) gene, which encodes the heme oxygenase that catalyses the conversion of heme to biliverdin IXa, the rate-limiting step in the synthesis of the phytochrome chromophore (Davis et al., 1999). Two independent Mp*hy1* mutants developed hyponastic thalli, while the wild types were flat (Figure 4B, C, Suppl. Figure 4C).

Collectively, these data indicate that red light perceived by phytochrome inhibits the development of thallus hyponasty. The data are consistent with the hypothesis that phytochrome-mediated red light signalling and phototropin-mediated blue light signalling antagonistically modulate thallus growth tropism.

### Mp*BBX* gene expression is induced by light before growth initiation (Figure 5)

To identify genes involved in light-modulated thallus tropism, we analysed transcriptional changes in growing thalli within 24 hours after exposure to light of different wavelengths. Thalli were grown for 8 days in white light and transferred to darkness for 24 hours. Plants were then transferred to white, blue, red, far-red light, or darkness and their time-resolved gene expression profiled (Figure 5A).

**Figure 5.**
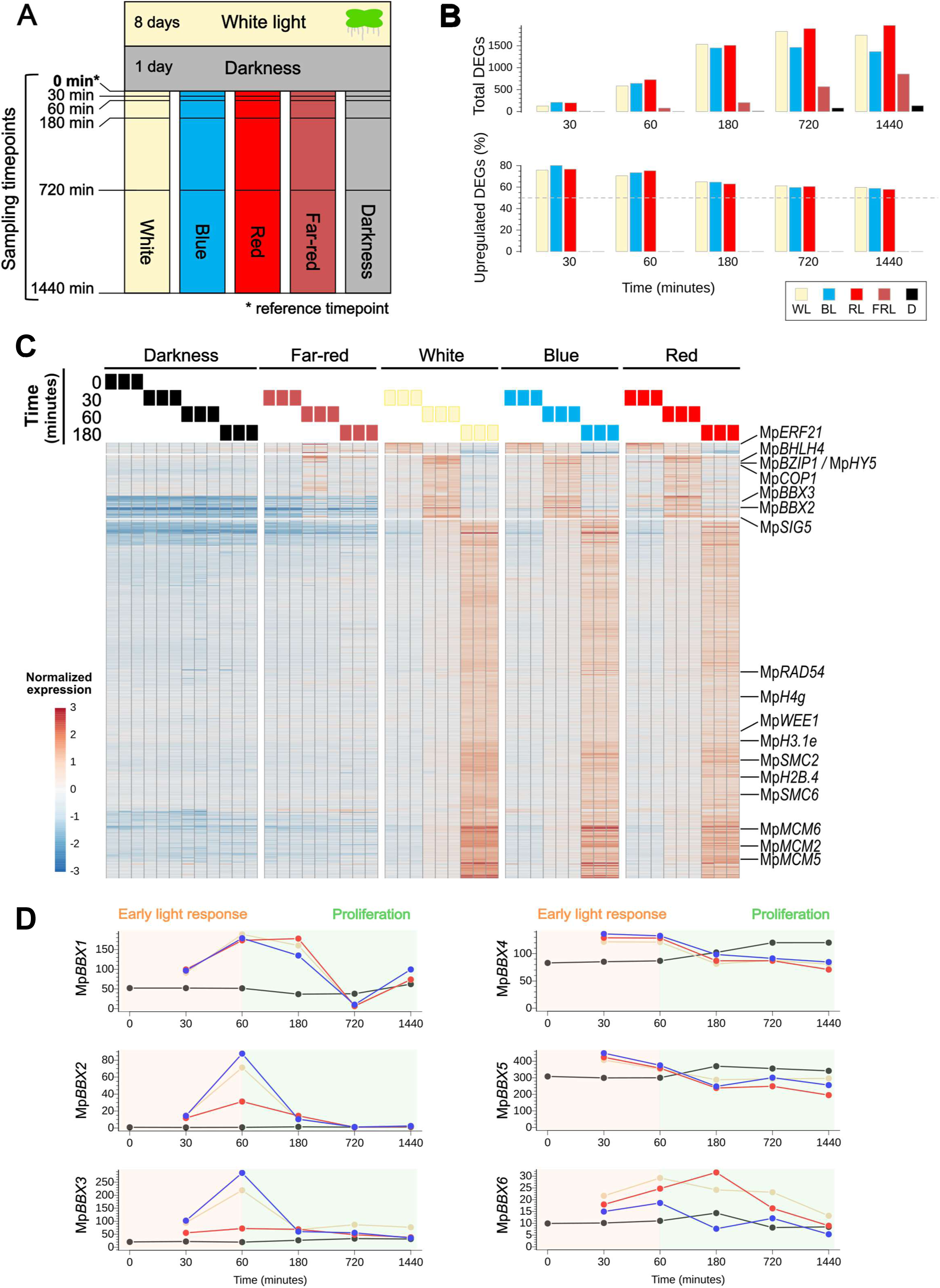
Wavelength-specific transcriptomes reveal that Mp*BBX* genes are light-induced. (A) Experimental design. Light treatments and sampling timepoints are shown. Plants were grown for 8 days in continuous white light, then transferred to complete darkness for one day. After the darkness incubation, they were transferred to their respective treatment at various wavelengths (darkness, white light, blue light, red light, far-red light) and harvested after 0, 30, 60, 180, 720, and 1440 minutes. (B) Differential gene expression analysis. Comparison of each transcriptome to the initial state (darkness timepoint 0) revealed significant transcriptional changes after 12 hours (720 minutes) and 24 hours (1440 minutes) of continuous light exposure. The first panel shows the total number of differentially expressed genes (DEGs), while the second panel illustrates the fraction of upregulated genes over time. The dashed line in the second panel represents the 50% threshold, where 50% depicts no net upregulation or downregulation. This data suggests an initial regulatory shift followed by broader transcriptional changes over time. (C) Clustered heatmap showing early transcriptional responses to light. Normalized gene expression is shown for all light conditions at timepoints 0 (reference), 30, 60, and 180 minutes. Selected genes are highlighted as representatives: Mp*ERF21*, induced within 30 minutes of light exposure; a set of light-responsive genes including Mp*BHLH4*, Mp*HY5*, Mp*COP1*, Mp*BBX3*, Mp*BBX2*, and Mp*SIG5* showing induction within 60 minutes; and a selection of genes encoding cell-cycle regulators that are light-induced within 180 minutes. (D) Light-induced expression of Mp*BBX* genes. Line plots demonstrate that all six *M. polymorpha BBX* genes were transcriptionally induced within 60 minutes of light exposure. Each plot represents the expression of the specific Mp*BBX* gene in response to darkness (black), white light (yellow), blue light (blue), and red light (red). The x-axis shows time in minutes, while the y-axis indicates normalized expression levels for each Mp*BBX* gene.

Comparison of each transcriptome to the initial state revealed extensive transcriptional changes after 12 hours (565-1889 differentially expressed genes (DEGs)) and 24 hours (854-1961 DEGs) of continuous light exposure (Figure 5B, Suppl. Figure 5 A). DEGs identified during the early timepoints (30-180 minutes) were mostly upregulated (Figure 5B). The fraction of upregulated genes decreased over time, suggesting gene expression changes between later timepoints (720 minutes and 1440 minutes) may be indirect downstream impacts of light rather than specific photo-regulation (Figure 5B). This was further supported by the high degree of overlap in gene expression in plants exposed for 12-24 hours of continuous white, blue or red light but not far-red light (only six differentially expressed genes (DEGs) were detected 30 minutes after transfer to far-red light (FRL) compared to the initial, dark state (timepoint 0; Suppl. Figure 5A, B). Early transcriptional responses to light, within 180 minutes, included a set of genes encoding S-phase cell-cycle regulators (Figure 5C), consistent with the known role of light in triggering growth-related pathways in plants (Casal & Yanovsky, 2005).

We hypothesized that the subtle expression changes observed by 30 minutes (6-207 DEGs), and 60 minutes (77-726 DEGs) would include genes whose early activation underpins light-modulated transcriptional reprogramming (Suppl. Figure 5A). All six *M. polymorpha BBX* genes – BBX transcription factors that act downstream of photoreceptors in land plants (Gangappa & Botto, 2014) – were transcriptionally induced within 60 minutes of light exposure (Figure 5D). This suggests that some of the BBX transcription factors may modulate early transcriptional changes that guide light-oriented thallus tropic growth. In blue-light, genes encoding canonical light-signalling regulators (including Mp*COP1*, Mp*RUP*, Mp*HY5*, Mp*SIG5*) were induced (Figure 5C, Suppl. Figure 5 C). In red light, Mp*BHLH4* expression was significantly induced after 30 and 60 min light treatment (Figure 5C, Suppl. Figure 5 C). Taken together, these data indicate that in *M. polymorpha* the expression of the Mp*BBX* gene family is induced by light. This supports the hypothesis that Mp*BBX* genes are involved in light-regulated development in *M. polymorpha*.

### Mp*BBX1* and Mp*BBX5* act antagonistically in thallus tropism (Figure 6)

BBX proteins are small transcription factors involved in light-regulated responses regulated by plant photoreceptors (Yadav et al., 2020). There are five structural groups of BBX proteins in land plants. There are six *BBX* genes in *M. polymorpha* in four of the five BBX structural groups (Suppl. Figure 6A). To test if Mp*BBX* genes encode proteins involved in thallus tropism, we generated mutants in each of the six Mp*BBX* genes using CRISPR-Cas9 mutagenesis (Figure 6A, Suppl. Figure 6B) and characterized the thallus growth tropism phenotype. Mp*bbx2,* Mp*bbx3,* Mp*bbx4,* and Mp*bbx6* mutants developed thallus phenotypes that were indistinguishable from wild type (Suppl. Figure 6C). These mutants displayed growth angles similar to those of F1 plants derived from a cross between Tak-1 and Tak-2, which fell within the range observed in the parent Tak-1 and Tak-2 plants (Suppl. Figure 6D). By contrast, thallus growth tropism in Mp*bbx1* and Mp*bbx5* was defective (Figure 6B, D). Mp*bbx1* mutants were hyponastic, resembling Mp*phy* defective mutants (Figure 6B, C). Mp*bbx5* mutants were epinastic, resembling Mp*phot* mutants (Figure 6D) with lower thallus curling indices (average 0.33) than wild type Tak-1 (average 0.93) (Figure 6E).

**Figure 6.**
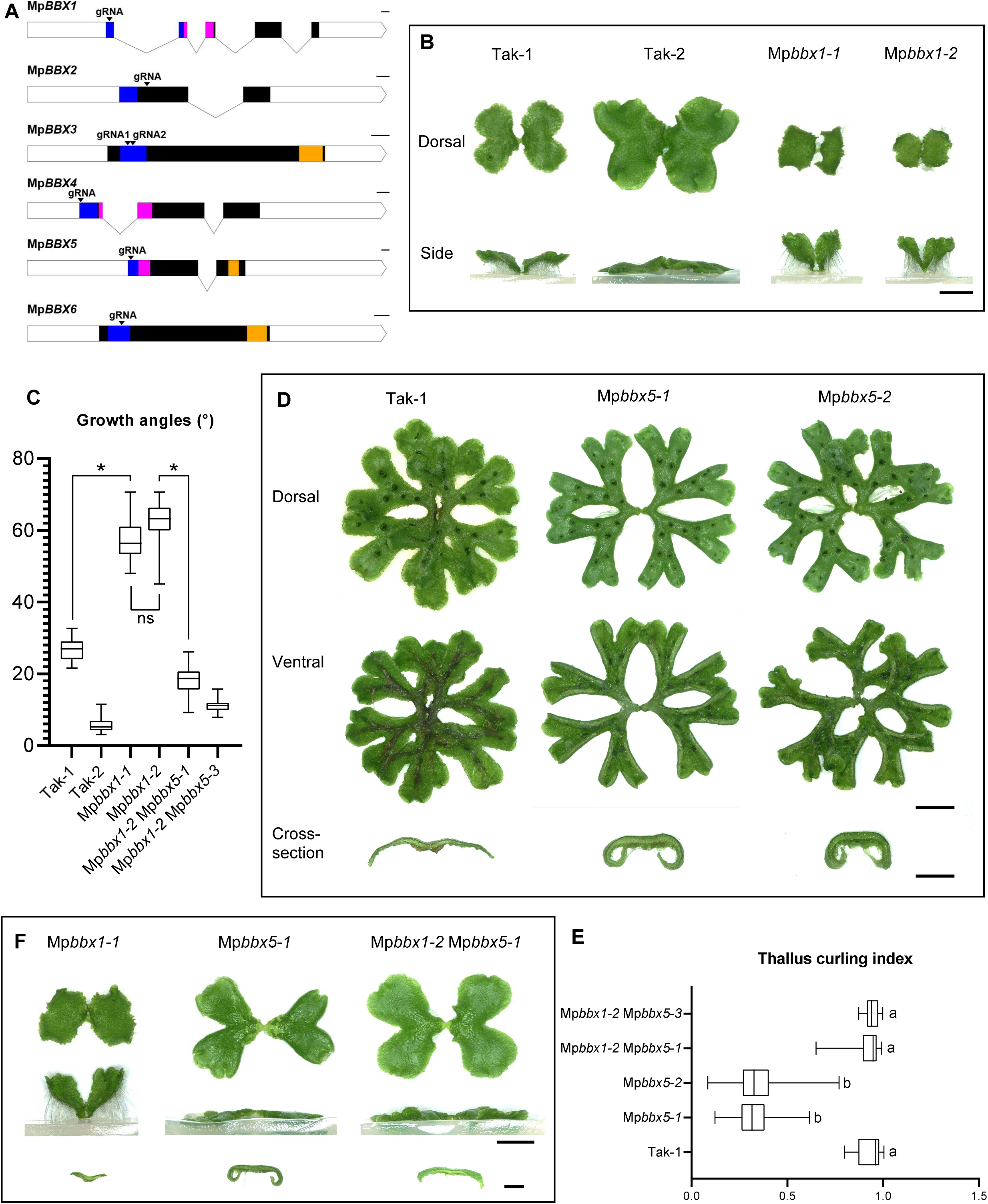
MpBBX1 is required for epinastic growth, and MpBBX5 is required for hyponastic growth. (A) CRISPR mutants for all six Mp*BBX* genes: sgRNA target sites are indicated with arrows in the gene models. The 5′ and 3′ UTRs are shown as white boxes, exons as black boxes connected by lines representing introns. Functional domains are color-coded: B-box1 in blue, B-box2 in pink, and the CCT domain in orange. Scale bar = 100 bp. (B) Detailed phenotype of Mp*bbx1* mutants showing hyponastic growth. Plants were grown for 14 days in continuous white light. Scale bar = 5 mm. (C) Quantification of growth angles in Mp*bbx1* mutants and Mp*bbx1* Mp*bbx5* double mutants compared to wild types (n = 95). Data were tested for normality using the Shapiro-Wilk test and for homogeneity of variances using Brown-Forsythe test. A Welch’s ANOVA was performed, followed by Dunnett’s T3 post-hoc test. Asterisks (*) in the comparisons denote statistically significant differences between groups (p < 0.05), while ‘ns’ indicates no significant difference (p > 0.05). (D) Detailed phenotype of Mp*bbx5* mutants showing epinastic growth. Plants were grown for four weeks on cellophane-covered medium in continuous white light. Scale bar = 10 mm; cross-section scale bar = 3 mm. (E) Thallus Curling Index (TCI) measurements for Mp*bbx5* mutants, Mp*bbx1* Mp*bbx5* double mutants and wild type. Data were tested for normality using the Shapiro-Wilk test and for homogeneity of variances using Brown-Forsythe test. A Welch’s ANOVA was performed, followed by Dunnett’s T3 post-hoc test. Groups with different letters are significantly different (p < 0.05), while identical letters indicate no significant difference. (F) The Mp*bbx1* Mp*bbx5* double mutants displayed a flat growth phenotype, indicating that MpBBX1 and MpBBX5 act antagonistically in controlling thallus tropism. Top and side views are from plants grown for 18 days in continuous white light; cross-sections were taken after three weeks. Scale bar = 5 mm for top and side views, 2 mm for cross-sections.

The hyponastic phenotype of Mp*bbx1* and the epinastic phenotype of Mp*bbx5* suggest that *BBX1* and *BBX5* antagonistically regulate thallus growth tropism. To test this hypothesis, we generated Mp*bbx1* Mp*bbx5* double mutants and determined the thallus growth tropism phenotype. While Mp*bbx1* mutants developed a hyponastic thallus and Mp*bbx5* mutants developed an epinastic thallus, the Mp*bbx1* Mp*bbx5* double mutants developed a flat thallus, resembling the wild type thallus growth tropism (Figure 6C, Figure6E-F). The phenotype of the Mp*bbx1* Mp*bbx5* double is consistent with the hypothesis that Mp*BBX1* and Mp*BBX5* act antagonistically in controlling thallus growth tropism. Together, these data indicate that Mp*BBX1* is required for epinastic growth and Mp*BBX5* is required for hyponastic growth and that these genes act antagonistically.

## Discussion

Light modulates the directional growth – tropism – of the *M. polymorpha* thallus, a system of dorsiventralized, indeterminate axes. Thalli respond to red and blue light in opposing ways: red light promotes downward tropism (epinasty) and blue light promotes upward tropism (hyponasty). In white light, the red and blue light activities balance each other, resulting in an intermediate, flat thallus (Figure 1). Although thalli are indeterminate axes and leaves are determinate, the antagonism of red-and blue light-controlled thallus tropisms we describe is similar to those described in *Arabidopsis* leaves (Kozuka et al., 2013). This suggests a deeply conserved photosensory mechanism across land plants, which orients flat dorsiventralized organs that are not homologous and have different developmental histories. We also show that the same photoreceptors function in blue and red light-modulated thallus tropism in *M. polymorpha* and leaf tropism in *A. thaliana.* This supports the hypothesis that the mechanism controlling the tropisms of flattened dorsiventralized structures is conserved among land plants. The blue light receptor PHOTOTROPIN is required for hyponastic growth – Mp*phot* mutants develop epinastic phenotypes in white light while wild type grows flat (Figure 2). We further show that the role of *M. polymorpha* PHOTOTROPIN in flattening tissues is conserved between *M. polymorpha* and *A. thaliana* and that its kinase activity is essential for this activity (Figure 3). The red light receptor PHYTOCHROME is required for epinastic growth – Mp*phy* mutants and Mp*hy1* mutants defective in the biosynthesis of the phytochrome chromophore, grow hyponastically while wild type thalli are flat (Figure 4).

To identify genes acting in red light- and blue light-modulated tropisms, we generated time-resolved transcriptomes from plants transferred from darkness into red or blue light (Figure 5). As expected, the mRNA levels of genes encoding regulators of canonical light signalling increased rapidly after exposure to blue light. Among these were *HY5*, *RUP* and *COP1*. Furthermore, *SIG5* is induced in our transcriptome within 60 minutes and has been previously shown to be required specifically for blue light induction of *psbD* transcription (Tsunoyama et al., 2002, Tsunoyama et al., 2004). Also induced within 60 minutes of blue light were transcripts corresponding to each of the six *BBX* genes encoded in the *M. polymorpha* genome. BBX proteins are transcriptional integrators of photoreceptor-mediated light signals and some BBX proteins have been shown to localize to photobodies in *A. thaliana* (Gangappa & Botto, 2014, Yadav et al., 2020, Cao et al., 2023, Van Buskirk et al., 2012). We demonstrate that of the six BBX genes in M. polymorpha, Mp*BBX1* and Mp*BBX5* are required for epinastic and hyponastic growth, respectively (Figure 6). Furthermore, they control tropism antagonistically; while the Mp*bbx1* single mutant is hyponastic and the Mp*bbx5* mutant is epinastic, the Mp*bbx1* Mp*bbx5* double mutant grows flat. We hypothesize that blue light-induced hyponasty is negatively regulated by Mp*BBX1* and positively regulated by Mp*BBX5*.

Fewer genes encoding proteins involved in light signalling were specifically induced by red light in our conditions than by blue light. However, both Mp*BBX1* and Mp*BBX5* were induced by red light and blue light consistent with the hypothesis that these regulators of thallus tropism are themselves controlled by red light as well as blue light. *bHLH4* transcript levels increase considerably by red light. We hypothesize that *bHLH4* is involved in red light transcriptional responses. There is no evidence that *bHLH4* or its orthologs are involved in tropisms in any plant. However, *bHLH4* is co-expressed with a group of genes hypothesized to be involved in developmental totipotency of *M. polymorpha* thalli, although there is no functional data to support this hypothesis (Flores-Sandoval et al., 2018).

While MpBBX1 and MpBBX5 are clearly involved in light-mediated growth responses in *M. polymorpha*, it is currently unknown if *A. thaliana* BBX proteins perform similar functions in tropic growth of leaves. MpBBX1 belongs to BBX group IV, and MpBBX5 to BBX group I. If the role of BBX proteins in regulating tropic growth is conserved across land plants, we predict that one or some group IV members (AtBBX18, AtBBX19, AtBBX20, AtBBX21, AtBBX22, AtBBX23, AtBBX24 and AtBBX25) would be required for epinastic leaf tropism. Similarly, we predict that group I members (AtBBX1, AtBBX2, AtBBX3, AtBBX4, AtBBX5, AtBBX6) would be required for hyponastic leaf tropism. If group IV and group 1 BBX proteins in *A. thaliana* control leaf tropism, it would be consistent with the hypothesis that components of the signalling cascade downstream of phototropin and phytochrome were also conserved among land plants.

The time-resolved, wavelength-specific transcriptomic dataset for *Marchantia polymorpha* thalli, which tracks gene expression dynamics under distinct light qualities (blue, red, far-red, and darkness) across multiple timepoints, represents the first comprehensive resource of its kind in bryophytes. As such, it offers a valuable tool for investigating light quality-dependent gene regulation in early-diverging land plants. To our knowledge, no comparable dataset exists for bryophytes. In this study, we developed an RShiny website (<u>mplight.dolan.vbc.ac.at</u> ; *Comment: the final version will run on a stable server*) to facilitate easy access to our time-resolved, wavelength-specific transcriptomic data, enabling users to explore individual *M. polymorpha* genes and their expression patterns under different light conditions and over time. This resource has the potential to facilitate the discovery of additional candidate genes involved in light signalling and photomorphogenesis.

## Conclusion

We conclude that in white light, flat thallus growth is maintained by the balance between the antagonistic activities of red light, which promotes epinastic growth, and blue light, which promotes hyponastic growth. Mp*PHY* and Mp*BBX1* are required for red light-induced epinastic growth, while Mp*PHOT* and Mp*BBX5* are required for blue light-induced hyponastic growth (Figure 7). Evidence supporting the role of these genes in the antagonistic modulation of tropisms are that the phenotypes of Mp*phy* and Mp*bbx1* loss-of-function mutants resemble wild type plants grown in blue light, while the phenotypes of Mp*phot* and Mp*bbx5* resemble wild type plants grown in red light. Furthermore, while the thallus of the Mp*bbx1* mutant is hyponastic and that of the Mp*bbx5* mutant is epinastic, the thallus of the Mp*bbx1* Mp*bbx5* double mutant is flat.

**Figure 7.**
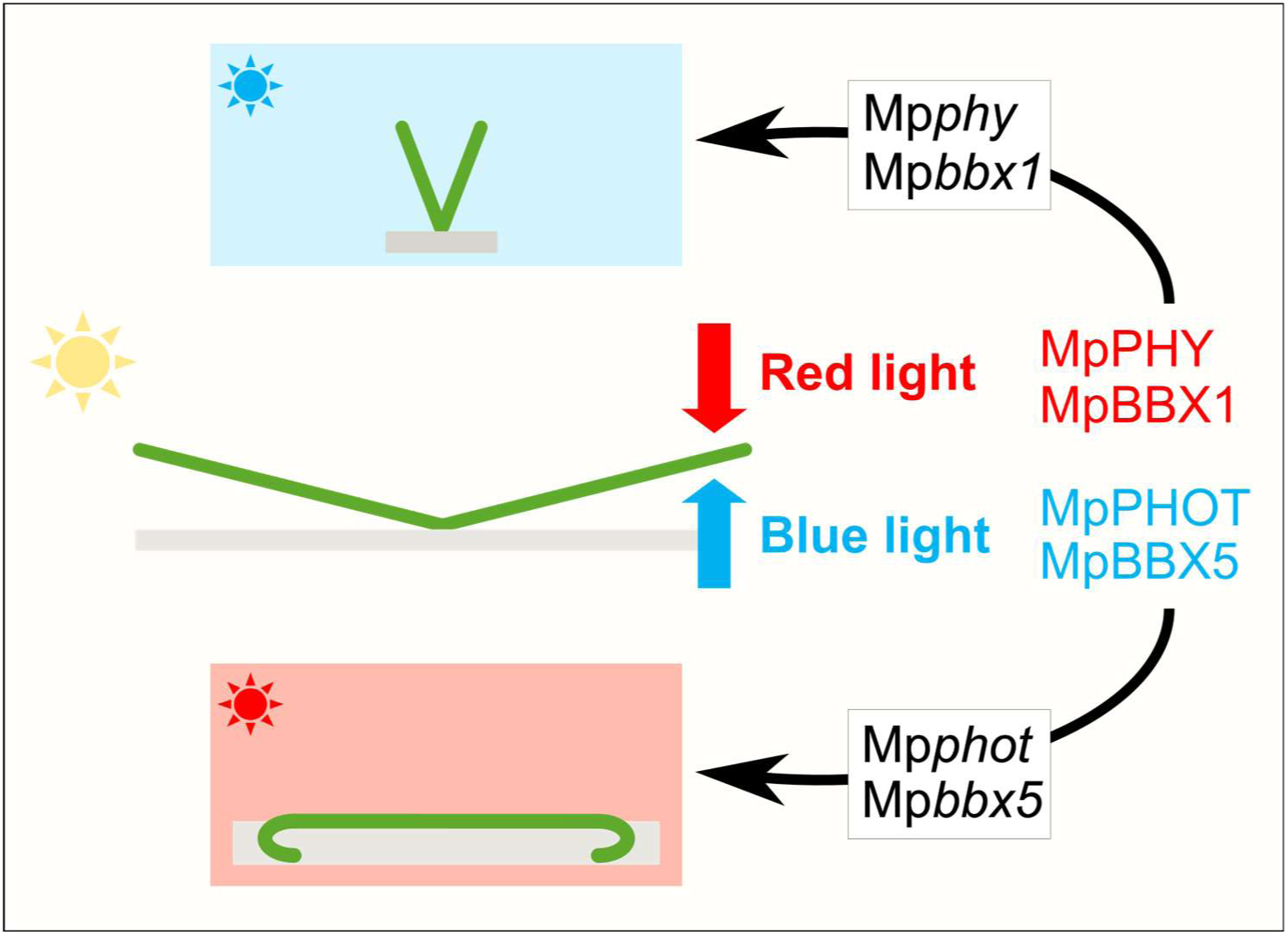
Schematic representation of the control of thallus tropism by red and blue light. In white light, flat thallus growth is maintained by the antagonistic functions of red light-promoted epinastic growth and blue light-promoted hyponastic growth. Mp*PHOT* and Mp*BBX5* promote hyponastic thallus tropism in blue light, and loss-of-function mutations in each gene result in an epinastic thallus tropism phenotype. Mp*PHY* and Mp*BBX1* promote hyponastic thallus tropism in red light and loss-of-function mutations in each gene result in a hyponastic thallus tropism phenotype.

## Acknowledgements

We are grateful to Magdalena Mosiolek and Katharina Jandrasits for general laboratory support. We thank John Christie and Stuart Sullivan for providing *Arabidopsis thaliana phot1-5 phot2-1* mutant seeds used for complementation experiments; Marco Incarbone for providing the GreenGate plasmid containing the selection marker for transformants (At*ATS3* seedcoat-specific promoter tagged to the fluorophore mCherry); Julia Kober for help and discussions regarding identification of target residue for generation of kinase-dead MpPHOT version; Eva-Sophie-Wallner for providing several general GreenGate plasmids (e.g., dummy modules) and the AtUBI10 promoter module; the ProTech Facility of Vienna BioCenter for cloning of Mp*PHY* overexpression constructs, Mathias Madalinksi (Peptide Synthesis Facility at the Vienna BioCenter) for synthesis of PHOT peptides and other core facilities at the Vienna BioCenter for constant support: BioOptics Facility, Plant Sciences Facility, the Media Lab, Lab Support and the administrative personnel at the VBC. We thank Thea E. Kongsted and Eva-Sophie Wallner for critically reading the manuscript. This study was funded by a grant from the Austrian Academy of Sciences (OEAW) to the Gregor Mendel Institute and a European Research Council advanced grant DENOVO-P (project number: 787613) to Liam Dolan.

## Materials and Methods

### Plant material and growth conditions

*Marchantia polymorpha* wild type accessions, Takaragaike-1 (Tak-1, male) and Takaragaike-2 (Tak-2, female), and mutants were grown on sterile plates with standard culture medium, half-strength B5 Gamborg with 0.5 g/L MES hydrate, 1% sucrose, and 1% plant agar, adjusted to pH 5.5.

Plants were grown in continuous light (see intensities in the respective figure legends) in growth chambers from poly klima Climatic Grow Systems (Hofmann Kühlung Company; firmware: 03.10.10(22), controller: WAGO 750-8212 PFC200 G2 2ETH R) at 23 °C. Darkness control sample plates were wrapped twice in aluminum foil, placed in a light-proof box, and kept in the plant growth chamber.

The *Arabidopsis thaliana* wild type accession, Columbia Col-0, and mutants and complementation lines were grown on soil under a 16-h light/8-h dark regime at 21°C. Whole plants and leaves were imaged after 25 days.

The following intensities were used if not indicated differently at the respective figures: Standard continuous white light conditions: PFD 50-60 μmol m⁻² s⁻¹; blue light-grown plants were exposed to ∼30-40 µmol m⁻² s⁻¹ blue light (λmax = 450 nm); red light-grown plants were exposed to ∼80-90 µmol m⁻² s⁻¹red light (λmax = 660 nm); far-red light-grown plants were exposed to 30-40 µmol m⁻² s⁻¹ far-red light (λmax = 730 nm); darkness controls were wrapped twice in aluminum foil, placed in a light-proof box, and kept in the plant growth chamber at 23°C. Light conditions were measured with the photometer Spectral PAR Meter PG100N (UPRtek). Screenshots from the different spectra and their λmax values are attached in Suppl. Figure 1 E.

### Growth angle measurements

For growth angle measurements, *M. polymorpha* plants were grown for 14 days under the specified light conditions, unless otherwise indicated in the corresponding figure. The agar surrounding a single plant was carefully cut into a rectangular shape and repositioned sideways, with the central connection point of the thallus placed at the center. The thallus lobes were oriented away from the center, as shown in all figures displaying growth angle measurement (Fig 1, 4, 5, 6).

After imaging the plants, the growth angle between the thallus tip and the agar surface was measured using the Angle Tool in FIJI and using the thallus tip, midpoint and agar surface as reference points. These measurements were taken for both sides of the thallus, resulting in two angle values for each individual.

### Thallus curling index

Cross-sections of the thallus were done by cutting single thallus branches of ∼3-week-old plants in a non-bifurcating area without gemmae cups in the middle of the branch into 1 mm thick pieces. The cross-section was placed onto a white background and imaged individually. For each genotype, several individual plants were used, and one cross-section was taken from each branch. To calculate the thallus curling index, the mid-point of the thallus cross-section and the tips of the thallus cross-section were used as reference points for the measurements (see Suppl. Fig 2).

Two lengths were measured for each side of the cross-section: 1) “real length”: the length between the midpoint and the tip of the thallus edge accurately measured along the ventral surface of the cross-section, and 2) “shortest distance”: the shortest distance between the midpoint and tip of the thallus edge measured by drawing a straight line between these two reference points. To calculate The Thallus Curling Index (TCI), these two values are divided: “shortest distance” / “real length” value. A TCI score of 1 describes a completely flat thallus. Length measurements were performed using the measurement software of the Keyence microscope VHX-7000.

### Leaf flattening index

*Arabidopsis thaliana* plants were grown on soil grown in long-day conditions (16-h light/8-h dark) for 25 days at 21°C. For each genotype, four plants and three leaves (L3, L4, L5) were used for calculating the leaf flattening index (LFI), resulting in 12 LFI measurements per genotype. Two images for each leaf were taken, one image of the unflattened abaxial leaf surface and one image of the artificially flattened leaf surface. To artificially flatten the leaf, cuts from the leaf margins into the leaf blade were performed and the leaf was then adhered onto a sticky tape with the adaxial side stuck onto the tape and the abaxial facing the camera as for the unflattened leaves. The surface area was defined with the measurement software of the Keyence microscope VHX-7000, manually adjusting the thresholds to define the leaf and background area. The LFI was calculated as the ratio of the surface of the unflattened and flattened leaf (unflattened: flattened). The flatter a leaf is the closer this value of the LFI is to 1. The definition of the leaf-flattening index as the ratio of curled to total projection areas (de Carbonnel et al., 2010) was adopted, with above-mentioned modifications made to the calculation based on the growth conditions, the number of leaves measured, and the use of the abaxial surface for measurements.

### CRISPR/Cas9 mutagenesis

#### sgRNA design and target site selection

CRISPR/Cas9-mediated mutagenesis was used to generate mutant lines for the following genes: Mp*BBX1* (Mp5g05940), Mp*BBX2* (Mp3g20050), Mp*BBX3* (Mp3g19670), Mp*BBX4* (Mp8g01190), Mp*BBX5* (Mp8g13470), Mp*BBX6* (Mp2g16350), Mp*HY1* (Mp6g10740), Mp*PHOT* (Mp5g03810). Single guide RNAs (sgRNAs) were designed using the CHOPCHOP online tool (Labun et al., 2019) or manually if the desired target region was not listed as one of the first hits. Gene sequences were retrieved from marchantia.info (Tanizawa et al., 2025), based on the MpTak v6.1r2 reference genome. Each sgRNA consisted of a 20-nucleotide target sequence followed by a canonical NGG PAM motif. To minimize off-target effects, each sgRNA candidate was screened via BLAST against the *M. polymorpha* genome to confirm unique targeting. Target sites, gRNA sequences, and the position of the gRNA relative to the whole gene sequence are indicated in the respective supplementary figures.

#### Generation of plasmids for transformation

To facilitate SapI-mediated loop assembly, overhang sequences were incorporated into primer designs. Forward primers were structured as TCG-[20 nt sgRNA]-GT, and reverse primers as AAAAC-[20 nt reverse complement of the sgRNA]. The resulting sgRNA duplexes were cloned into the OP-076 vector (L2_lacZgRNA-Cas9-CsA) from the OpenPlant kit, following the assembly method described by (Sauret-Güeto et al., 2020). The double gRNA construct for mutagenesis of MpHY1 was also cloned using the vectors from the Open Plant toolkit following the corresponding cloning protocol. The vectors carry dual antibiotic resistance markers for hygromycin and spectinomycin, enabling selection of plasmid-carrying transformants.

#### Bacterial transformation and plasmid sequencing

Recombinant plasmids were introduced into *Escherichia coli* DH5α cells via heat shock. Transformed colonies were screened on LB agar plates supplemented with 50 μg/mL spectinomycin and 40 μg/mL X-Gal for blue-white selection. White colonies were cultured overnight in liquid LB medium containing spectinomycin, and plasmids were extracted using the MiniPEx kit (Molecular Biology Service, Vienna BioCenter). The correct insertion of the sgRNA cassette into the CRISPR/Cas9 vector was verified by Sanger sequencing using the Mp-U6-seqF primer (GAGTTCGGATTGCTCTCTTTC), performed at the Vienna BioCenter core facility.

#### Generation of transgenic lines

Mutant lines were generated by spore transformation with spores generated from a cross of Tak-1 and Tak-2.

#### *Agrobacterium*-mediated transformation of *Marchantia polymorpha* sporelings

Purified plasmids were electroporated into *Agrobacterium tumefaciens* strain GV3101, which is resistant to rifampicin and gentamycin. Transformants were selected on LB agar plates containing rifampicin, gentamycin, and spectinomycin (each at 50 μg/mL), and incubated in darkness at 28 °C for two days. Individual colonies were subsequently cultured in liquid LB medium containing the same antibiotics. Transformation of *M. polymorpha* was carried out according to the protocol described by (Ishizaki et al., 2008), with minor modifications. Dried and frozen wild type sporangia, derived from a cross of the two wild type accessions Tak-1 and Tak-2, were sterilized in 0.1% sodium dichloroisocyanurate (NaDCC; Sigma-Aldrich, Cat. No. 218928), washed with sterile distilled water, and suspended in 1 mL dHCO. The resulting spore suspension was cultured in 25 mL M51C medium (Takenaka et al., 2000) under continuous white light at 23 °C for seven days.

*Agrobacterium* GV3101 strains harbouring the sgRNA constructs were grown for two days in darkness at 28 °C in 5 mL liquid LB medium containing 50 μg/mL of rifampicin, gentamycin, and spectinomycin. Bacterial cells were pelleted by centrifugation and resuspended in 10 mL M51C medium supplemented with 100 μM acetosyringone, followed by a 4-hour induction at 28 °C in the dark. One milliliter of the induced *Agrobacterium* culture was added to the sporeling culture and co-cultivated for two days in the presence of 100 μM acetosyringone. Following co-cultivation, the sporelings were repeatedly washed and plated onto selective medium containing 100 μg/mL cefotaxime and 50 μg/mL hygromycin. The plates were kept in continuous white light at 23 °C >2 weeks until transformants were bis enough to be transferred to fresh selective plates (Ishizaki et al., 2008).

#### Mutant screening and genotyping

Genomic DNA was extracted from ∼3x3 mm thallus tissue from up to 50 hygromycin-resistant transformants for each gRNA construct. Samples were homogenized in 100CμL of extraction buffer containing 100 mM Tris-HCl (pH 9.5), 1CM KCl, and 10CmM EDTA. The resulting lysates were incubated at 65 °C for 10-15 minutes, followed by a 1:5 dilution with sterile Mono Q water. 1 μL of the diluted genomic DNA was used as the template for the PCR. PCR amplification was carried out using a 2x in-house master mix (Molecular Biology Service, Vienna BioCenter), which includes a hot start DNA polymerase. Each reaction contained 0.5 μM of both forward and reverse primers. Thermocycling conditions consisted of 35 cycles of denaturation at 95 °C for 30 seconds, annealing at 55 °C for 30 seconds, and extension at 72 °C for 1 minute/kb DNA. The 20 μL PCR product was treated with 2CμL of ExoSAP purification mix (each containing of 0.04 µl exonuclease I, 0.4 µl shrimp alkaline phosphatase, and 1.56 µl storage solution - 10 mM Tris-HCl, 100CmM NaCl, 5 mM β-mercaptoethanol, 0.5 mM EDTA, 50% glycerol, 1CmM MgCl2, pH 7.5). The digestion reaction was performed at 37 °C for 30 minutes and then heat-inactivated at 80 °C for 10 minutes. For mutation screening, 1 μL of the cleaned PCR product was sent to Sanger sequencing. CRISPR mutants were identified by amplifying regions spanning the gRNA target sites in the Mp*BBX1*, Mp*BBX2*, Mp*BBX3*, Mp*BBX4*, Mp*BBX5*, Mp*BBX6*, Mp*HY1*, Mp*PHOT* genes. The sequences were aligned against the wild type genome sequence using the CLC Genomics Workbench to identify mutations at the target sites. Lines with mutations in the targeted regions resulting in premature stop codons were selected for experiments. Gemmae or thallus pieces derived from these lines were propagated and repeatedly genotyped to confirm the presence of mutations.

#### Generation of plasmids for cross-species complementation experiments

We followed the GreenGate protocol (Lampropoulos et al., 2013) to generate the complementation constructs for experiments with *A. thaliana* and to generate the translational reporter construct in *M. polymorpha*.

The regions 4671 bp (for AtPHOT1), 3047 bp (for AtPHOT2), and 2490 bp (for MpPHOT) upstream of the start codon were defined as promoters for the respective genes. The coding sequence of MpPHOT comprises 3345 bp (without the STOP codon). After PCR amplification of the respective sequences with the sticky ends according to the GreenGate protocol, they were cloned into the GreenGate entry modules (pUC19-based vector backbone) via BsaI restriction sites. To generate the final constructs, the entry modules were assembled into the GreenGate destination vector (pGreen-IIS based). The linker-VENUS module was previously published (Wallner et al., 2023), the chlorsulfuron plant resistance module was modified from the OpenPlant toolkit (Sauret-Güeto et al., 2020).

#### Generation of plasmids for the kinase inactive version of MpPHOT

To generate a kinase-inactive version of MpPHOT, the aspartic acid codon in the conserved YRD motif was mutated to encode asparagine (YRN), resulting in a D904N substitution. This amino acid change was introduced into a plasmid containing the MpPHOT coding sequence using the QuikChange II Site-Directed Mutagenesis Kit (Agilent, product number: 200523), following the manufacturer’s protocol. Primers for the mutagenesis were designed using Agilent’s online Primer Design Program (forward primer: CGTTTTCAGGTTTCAGGTTCCTGTATACAACACCCAT; reverse primer: ATGGGTGTTGTATACAGGAACCTGAAACCTGAAAACG). Following confirmation of the mutation by Sanger sequencing, the plasmid was used as a GreenGate entry module and assembled into the final constructs as described in Fig. 3.

#### *Arabidopsis thaliana* transformation and generation of homozygous transgenic lines

*Arabidopsis thaliana* plants were cultivated under long-day photoperiods (16 hours light / 8 hours dark) for five weeks. Shortly after bolting, the primary inflorescence stems were trimmed to encourage the development of multiple lateral branches, leading to a higher number of floral buds and thus improved transformation rates. Around 10 days after the initial cut, plants produced lateral branches with an abundance of open and closed flower buds, the optimal stage for floral dip transformation.

*Agrobacterium tumefaciens* colonies harbouring the desired constructs were grown in 5 mL of LB liquid medium supplemented with hygromycin, gentamycin, and rifampicin (each at 50 μg/mL). Cultures were incubated at 28 °C, shaking at 150Crpm for 2 days. Subsequently, 250 mL of fresh LB medium containing the same antibiotics was inoculated with the starter culture and incubated under identical conditions for another 2 days.

The transformation procedure followed the floral dip method described by (Clough & Bent, 1998), with minor modifications. The 250 mL bacterial culture was centrifuged to pellet the cells, which were then resuspended in 500 mL of a 5% (w/v) sucrose solution. To facilitate *Agrobacterium* penetration into plant tissue, 0.02% Silwet L-77 was added to the suspension.

Plants were immersed in the *Agrobacterium* suspension for about two minutes while being gently swirled. Following dipping, plants were put into plastic bags to maintain high humidity and incubated overnight in the dark to enhance transformation efficiency. The following day, plants were returned to growth chambers under long-day conditions. Seeds were collected approximately one month after transformation.

Transgenic seeds were identified based on a fluorescent marker encoded by the F module of the GreenGate system. Seeds carrying the transgene exhibited red fluorescence in their seed coats (mCherry), which was visualized under a binocular microscope equipped with green laser illumination.

Fluorescent seeds were then surface-sterilized and subjected to cold stratification at 4C°C in darkness for two days. Afterwards, they were transferred to a growth chamber and grown under long-day conditions. Seeds collected from these independent transformants were re-screened for the presence of the transgene and selected accordingly. Another round of growth and selection was conducted to identify homozygous lines in independent lines for all constructs. These homozygous transgenic lines were used for experiments.

#### Protein extraction and western blotting

Plant material was harvested from 2-week-old plants grown on cellophane on standard medium in continuous white light and flash-frozen in liquid nitrogen. The frozen tissue was then ground to a fine powder using a liquid nitrogen-cooled mortar and pestle and transferred to a 1.5 mL Eppendorf tube filled to the 50 µL mark.

Total protein extraction was done by adding 100 µl of 5X Laemmli buffer (Laemmli buffer recipe for 15 ml total volume: 1.5 g SDS, 1M Tris pH=6.8, 0.015 g bromophenol blue, 1.16 g DTT, 7.5 ml glycerol, distilled water up to 15 mL total volume) to the ground tissue and vortexing rigorously. This step was followed by heating the samples at 95°C for 5 minutes, shaking at 300 rpm. The samples were then centrifuged at maximum speed for 5 minutes. 5 μL of the supernatant per sample, together with a protein ladder (Thermo Scientific Prestained Protein Ladder, 10 to 250 kD), were loaded into a precast protein gel (4-20% Mini-Protean TGX) in a Bio-Rad Mini-PROTEAN chamber filled with 1xSDS buffer. After running the gel, the separated proteins from the gel were electroblotted (Trans-Blot Turbo, 1.5 MM GEL program, Bio-Rad) onto a methanol-activated PVDF membrane. The membrane was blocked for 1 hour at room temperature using 5% skim milk in 1x TBS-T to prevent non-specific binding of the antibody. 20 µL of the custom-made anti-MpPHOT peptide antibody was added (1:1000) to the blocking solution. The rabbit antibody (anti-MpPHOT, primary antibody, custom-made by Eurogentec, Belgium) was raised against the peptide sequence (C+VDERAPPSKGSAKE) which was previously published (Fujii et al., 2017). The membrane was incubated with the primary antibody overnight at 4°C.

The membrane was washed three times with 1x TBS-T 20 min each. As a secondary antibody, 2 µL of the anti-rabbit IgG-HRP antibody (1:5000 dilution) in 5% Skim milk were added in 1xTBS-T buffer and incubated for 2 hours at room temperature. After washing the membrane three times with 1x TBS-T 20 min each, the detection solution SuperSignal™ West Pico PLUS Chemiluminescent Substrate (ThermoFisher) and the ChemiDoc Touch Imaging System 2385 (Bio-Rad) were used to detect proteins. Coomassie Blue staining served as a loading control.

#### Bright field plant imaging

Plants were imaged using the Keyence digital bright field microscope VHX-7000 equipped with the VH-Z00R/Z00T and VH-ZST lenses and the VHX-7020 camera if not otherwise indicated. If required, unspecific background from images was removed and images cropped to adequate size for figure assembly.

#### Surface area measurements

Thallus and gemmaling surface area were measured with the Keyence imaging software, adapting the selection of plant tissue with the extraction tools to define background and plant surface manually.

#### Confocal images of MpPHOT-VENUS localization in Marchantia polymorpha and Arabidopsis thaliana

Gemmae with the translational fusion construct *pro*Mp*PHOT:*Mp*PHOT-VENUS* were grown for 2 days in continuous white light (PFD 50-60 μmol m^-2^ s^-1^) prior to imaging. Images of the leaves from the complemented At*phot1*At*phot2* line, harbouring the *pro*At*PHOT2:*Mp*PHOT-VENUS*, were taken from plants grown on soil under a 16-h light/8-h dark regime at 21°C. Samples were imaged with an Olympus IX3 Series (IX83) inverted microscope, equipped with a Yokogawa W1 spinning disk, a Hamamatsu ORCA-Fusion CMOS camera and a 20x/0.8 or 40x/0.75 NA air objective. The samples were excited at 514 nm and emission was detected between 520-540 nm. The z-stacks were taken with 1-2 µm slices. Images were analyzed in ImageJ/FIJI.

#### Total RNA isolation, RNA sequencing and analysis

Wild type Tak-1 gemmae were transferred onto medium plates (½ Gamborg, MES, 1% sucrose, and 1% agar), each plate containing three biological replicates.

Gemmae were cultured under continuous white light at 23°C for eight days, followed by a 24-hour dark incubation. After this dark period, the plates were either kept in darkness (control) or exposed to different light conditions: broad white light (WL), monochromatic blue light (BL, λmax = 430 nm), monochromatic red light (RL, λmax = 660 nm), or monochromatic far-red light (FRL, λmax = 730 nm). The light intensity in the monochromatic light treatments was increased by a factor of four, relative to the intensity of the respective wavelength in the standard white light condition.

For each condition, three biological replicates - each containing three individual plants - were collected in 2 mL Eppendorf tubes containing a stainless-steel bead (5 mm) and immediately frozen in liquid nitrogen at the defined timepoints of 0 min, 30 min, 60 min, 180 min, 12 hours, and 24 hours in dark, white light, blue light, red, far-red light, respectively. Each sample consisted of two Tak-1 plants. All samples were stored at -70°C until the full set was collected. Tissue from the samples was disrupted using a mixer mill (RETSCH MM 400) for 4 minutes at 30 Hz, with nitrogen-based cooling breaks during the process. Total RNA was extracted using the High-Performance RNA Bead Isolation kit from the Molecular Biology Service facility at the Vienna BioCenter. The kit employs a guanidine thiocyanate-based lysis method adapted from (Boom et al., 1990) and utilizes carboxylate-modified Sera-Mag Speed beads, with processing done on the KingFisher instrument (Thermo).

mRNA library preparation, including poly(A) capture and NovaSeq S4 PE150 XP RNA sequencing, was performed by the Next Generation Sequencing (NGS) VBC Core Facility. The 78 cDNA libraries, corresponding to 26 treatments with three biological replicates each, were sequenced, yielding between 20.1 and 50.1 million reads per library (average of 38.4 million). Mapping and counting of transcripts were performed using the nf-core RNA-seq pipeline v3.14.0 (Di Tommaso et al., 2017, Harshil Patel, 2024) on the VBC CLIP Batch Environment (CBE) cluster. Raw reads were mapped to the *M. polymorpha* Tak-1 reference genome assembly MpTak1_v5.1 and its corresponding gene annotation version MpTak1_v5.1_r1 (marchantia.info) using STAR (Dobin et al., 2013). Reads were normalized by the sequencing depth of each library. After confirming low variability among biological replicates, the average expression value of each gene in each treatment was computed and used for downstream analyses. Differentially expressed genes (DEGs) were defined for each treatment as those with a logC fold change greater than 1 or less than –1 relative to the reference sample (0 hours).

#### Phylogenetic Analysis

The full-length amino acid sequences of the six BBX proteins from *Marchantia polymorpha* were obtained from marchantia.info (Tanizawa et al., 2025), based on the MpTak v6.1r2 reference genome, which comprises the male Tak-1 genome and the female Tak-2 U chromosome. The full-length amino acid sequences for the 32 BBX proteins from *Arabidopsis thaliana* were retrieved from The Arabidopsis Information Resource (Rhee et al., 2003).

These sequences, along with additional BBX protein sequences from twelve plant species (Crocco & Botto, 2013), were used for full-length amino acid sequence alignment. This set included sequences from four green algae, one moss, one lycophyte, three monocot and two dicot species as provided in the supplementary materials of the original publication (Crocco & Botto, 2013). In total, 220 protein sequences were aligned using MAFFT version 7 (Katoh et al., 2019) employing the L-INS-I method for iterative refinement, with all other parameters set to default.

Phylogenetic analysis was conducted using the maximum likelihood method implemented in IQ-TREE (Nguyen et al., 2015, Letunic & Bork, 2021), with default settings and ultrafast bootstrapping (Hoang et al., 2018). Model selection was performed using ModelFinder (Kalyaanamoorthy et al., 2017), which identified JTT-I-G4 as the best-fit model according to the Bayesian Information Criterion (BIC). An unrooted phylogenetic tree with bootstrap support values was visualized using iTOL version 6.9 (Letunic & Bork, 2021).

Motif analysis was carried out using MEME version 5.5.5 (Bailey & Elkan, 1994), accessed via the MEME Suite web server (Bailey et al., 2015), with default settings. Three conserved motifs were identified across the BBX proteins, with E-values ranging from 4.10 e–15 to 3.70 e–67. Nuclear localization signals (NLS) in MpBBX protein sequences were identified through manual screening and subsequently validated using LOCALIZER version 1.0.4 (Sperschneider et al., 2017).

#### Quantifications and statistical analysis

Sequence assemblies and plasmid maps were done in CLC Genomics Workbench 9.5.1 (Qiagen). The gene sequences were obtained from MarpolBase (https://marchantia.info/) (Tanizawa et al., 2025). All experiments were repeated 3-6 times. Sample sizes are indicated in the figure legends.

Analyses and plots were done in RStudio (version 2022.07.2 Build 576) or GraphPad Prism (version 8.1.1 (330); all statistical tests were performed in GraphPad Prism.

