## Supplementary material for "Antagonism between blue and red light-signalling controls thallus flatness in *Marchantia polymorpha*": Suppl. Figures

### Suppl. Figures 1-6

# S1

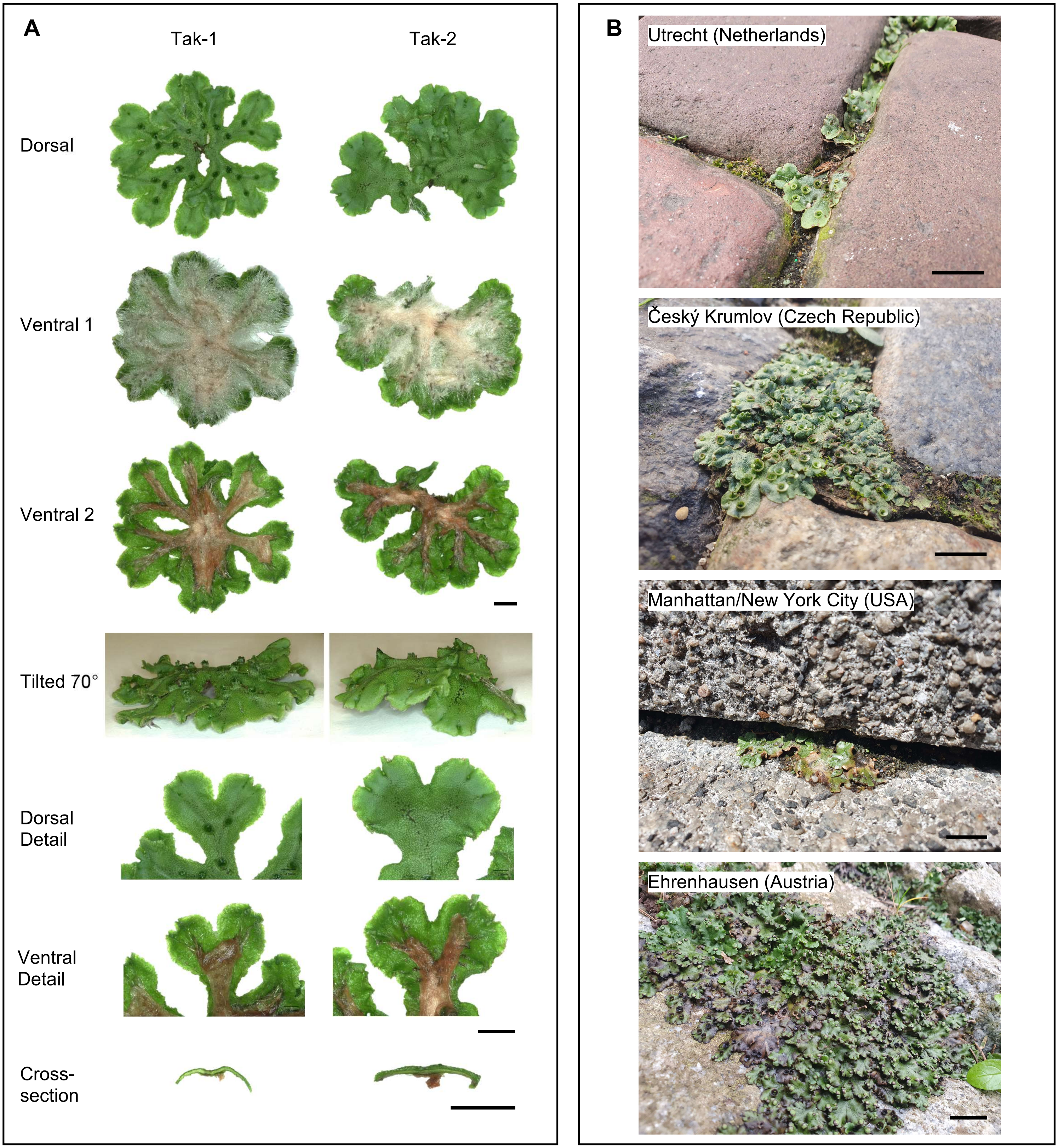
A
Tak-1
Tak-2
Dorsal
Ventral 1
Ventral 2
Tilted 70°
Dorsal Detail
Ventral Detail
Cross-section
B
Utrecht (Netherlands)
Český Krumlov (Czech Republic)
Manhattan/New York City (USA)
Ehrenhausen (Austria)

# S1

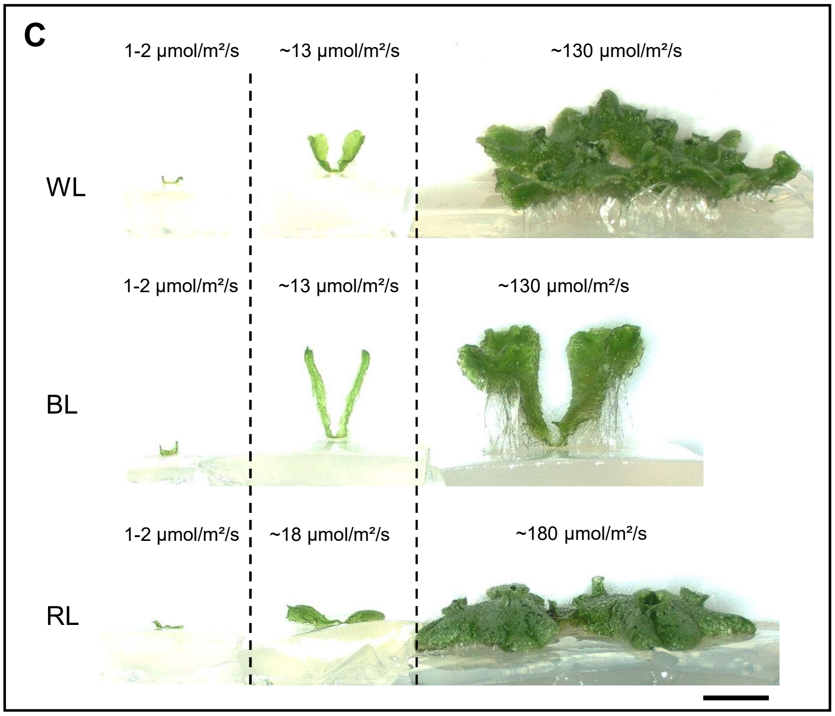
C
1-2 µmol/m²/s
~13 µmol/m²/s
~130 µmol/m²/s
WL
1-2 µmol/m²/s
~13 µmol/m²/s
~130 µmol/m²/s
BL
1-2 µmol/m²/s
~18 µmol/m²/s
~180 µmol/m²/s
RL

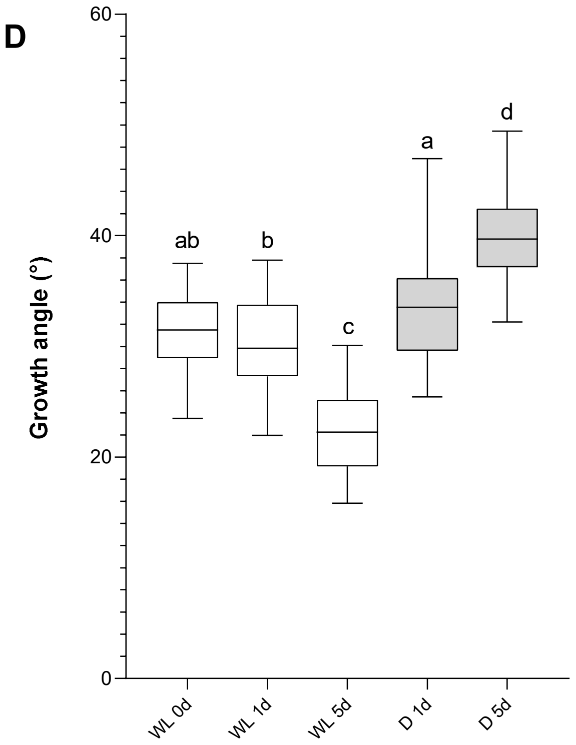
D
Growth angle (°)
60
40
20
0
ab
b
c
a
d
WL 0d
WL 1d
WL 5d
D 1d
D 5d

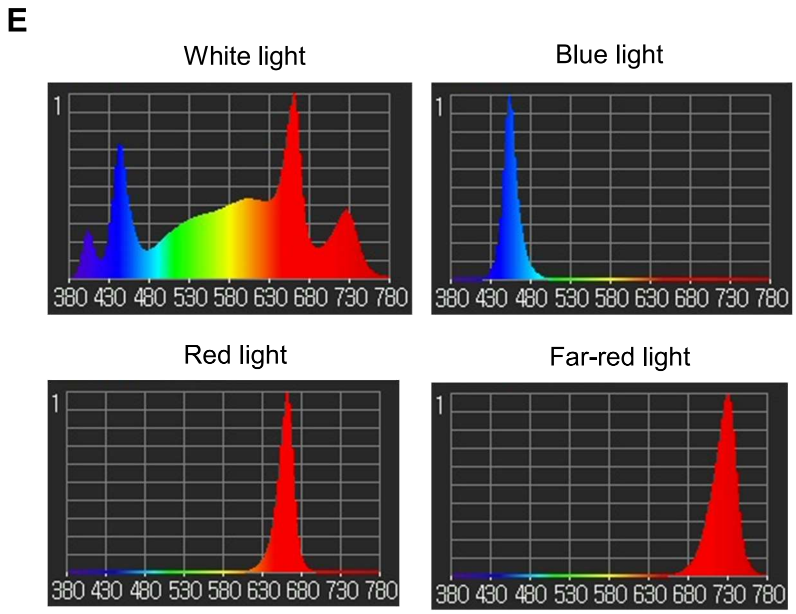
E
White light
Blue light
Red light
Far-red light
1
380 430 480 530 580 630 680 730 780

S2

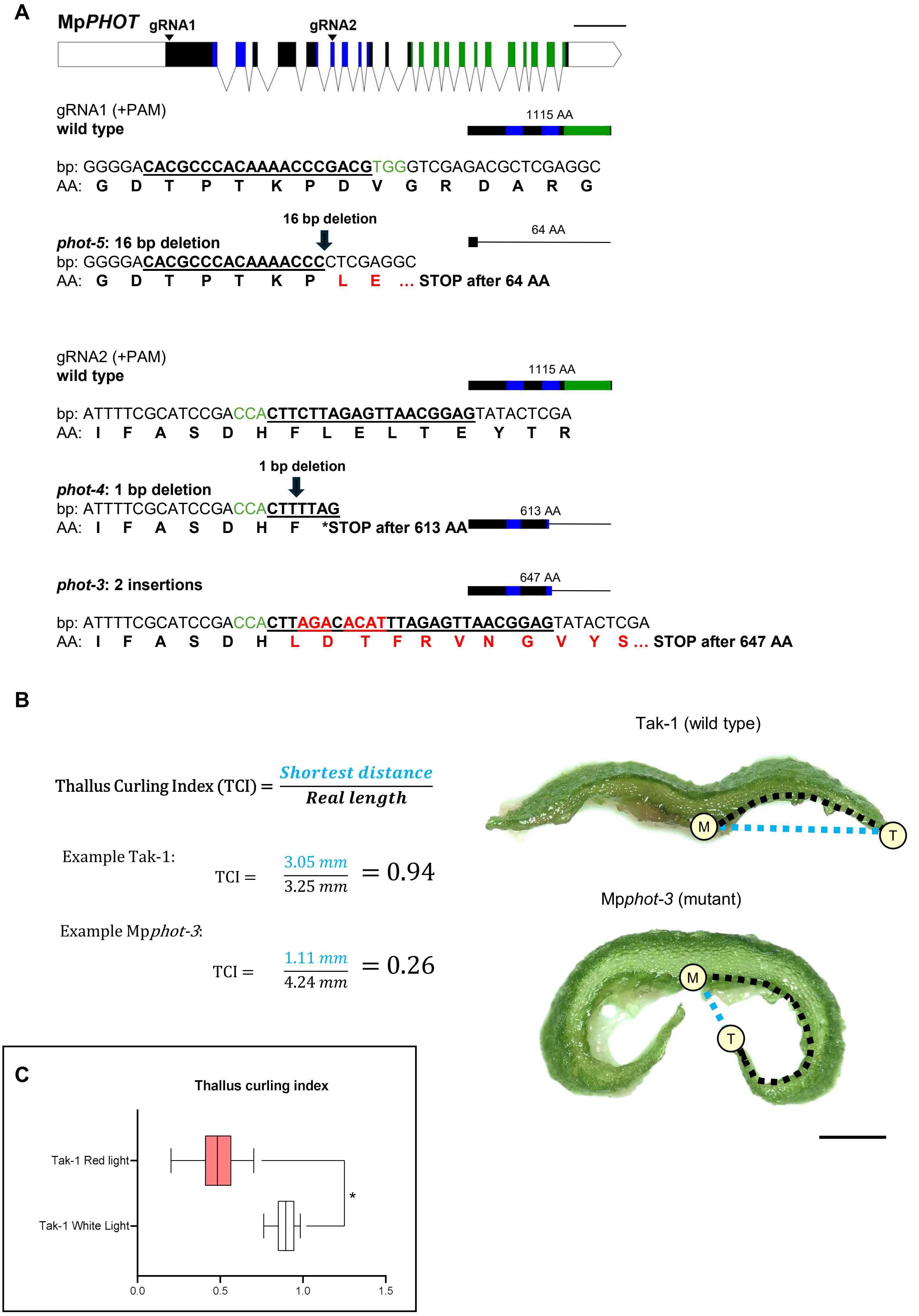
A
MpPHOT
gRNA1
gRNA2
gRNA1 (+PAM)
wild type
1115 AA
bp: GGGGACACGCCCACAAAACCCGACGTGGGTCGAGACGCTCGAGGC
AA: G D T P T K P D V G R D A R G
16 bp deletion
phot-5: 16 bp deletion
64 AA
bp: GGGGACACGCCCACAAAACCCCTCGAGGC
AA: G D T P T K P L E ... STOP after 64 AA
gRNA2 (+PAM)
wild type
1115 AA
bp: ATTTTCGCATCCGACCACTTCTTAGAGTTAACGGAGTATACTCGA
AA: I F A S D H F L E L T E Y T R
1 bp deletion
phot-4: 1 bp deletion
bp: ATTTTCGCATCCGACCACTTTTAG
AA: I F A S D H F *STOP after 613 AA
613 AA
phot-3: 2 insertions
647 AA
bp: ATTTTCGCATCCGACCACTTAGACACATTTAGAGTTAACGGAGTATACTCGA
AA: I F A S D H L D T F R V N G V Y S ... STOP after 647 AA
B
Thallus Curling Index (TCI) = Shortest distance / Real length
Example Tak-1:
TCI = 3.05 mm / 3.25 mm = 0.94
Example Mpphot-3:
TCI = 1.11 mm / 4.24 mm = 0.26
Tak-1 (wild type)
M
T
Mpphot-3 (mutant)
M
T
C
Thallus curling index
Tak-1 Red light
Tak-1 White Light
*
0.0
0.5
1.0
1.5

S3

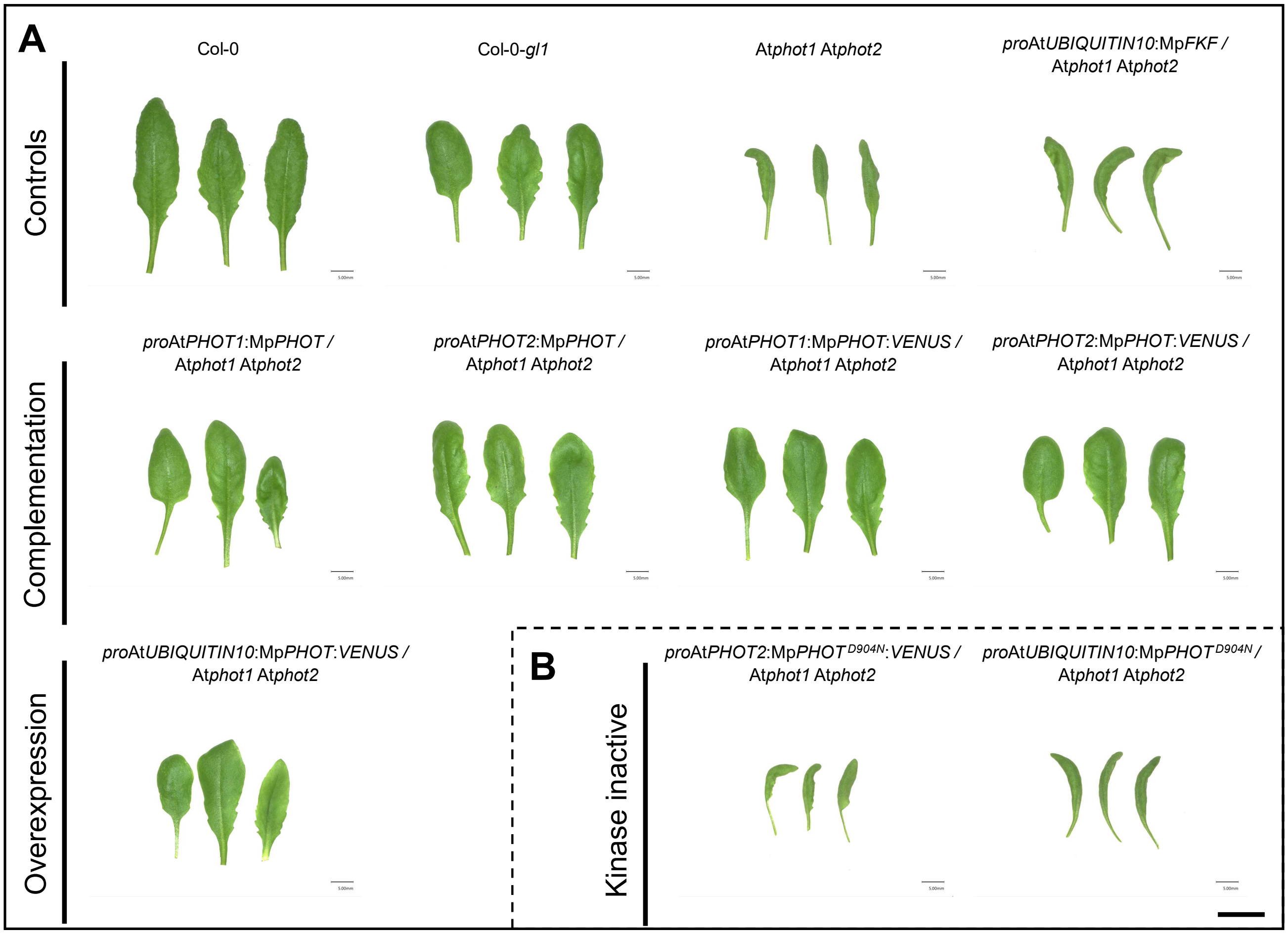
A
Controls
Col-0
Col-0-gl1
Atphot1 Atphot2
proAtUBIQUITIN10:MpFKF / Atphot1 Atphot2
Complementation
proAtPHOT1:MpPHOT / Atphot1 Atphot2
proAtPHOT2:MpPHOT / Atphot1 Atphot2
proAtPHOT1:MpPHOT:VENUS / Atphot1 Atphot2
proAtPHOT2:MpPHOT:VENUS / Atphot1 Atphot2
Overexpression
proAtUBIQUITIN10:MpPHOT:VENUS / Atphot1 Atphot2
B
Kinase inactive
proAtPHOT2:MpPHOT D904N:VENUS / Atphot1 Atphot2
proAtUBIQUITIN10:MpPHOT D904N / Atphot1 Atphot2

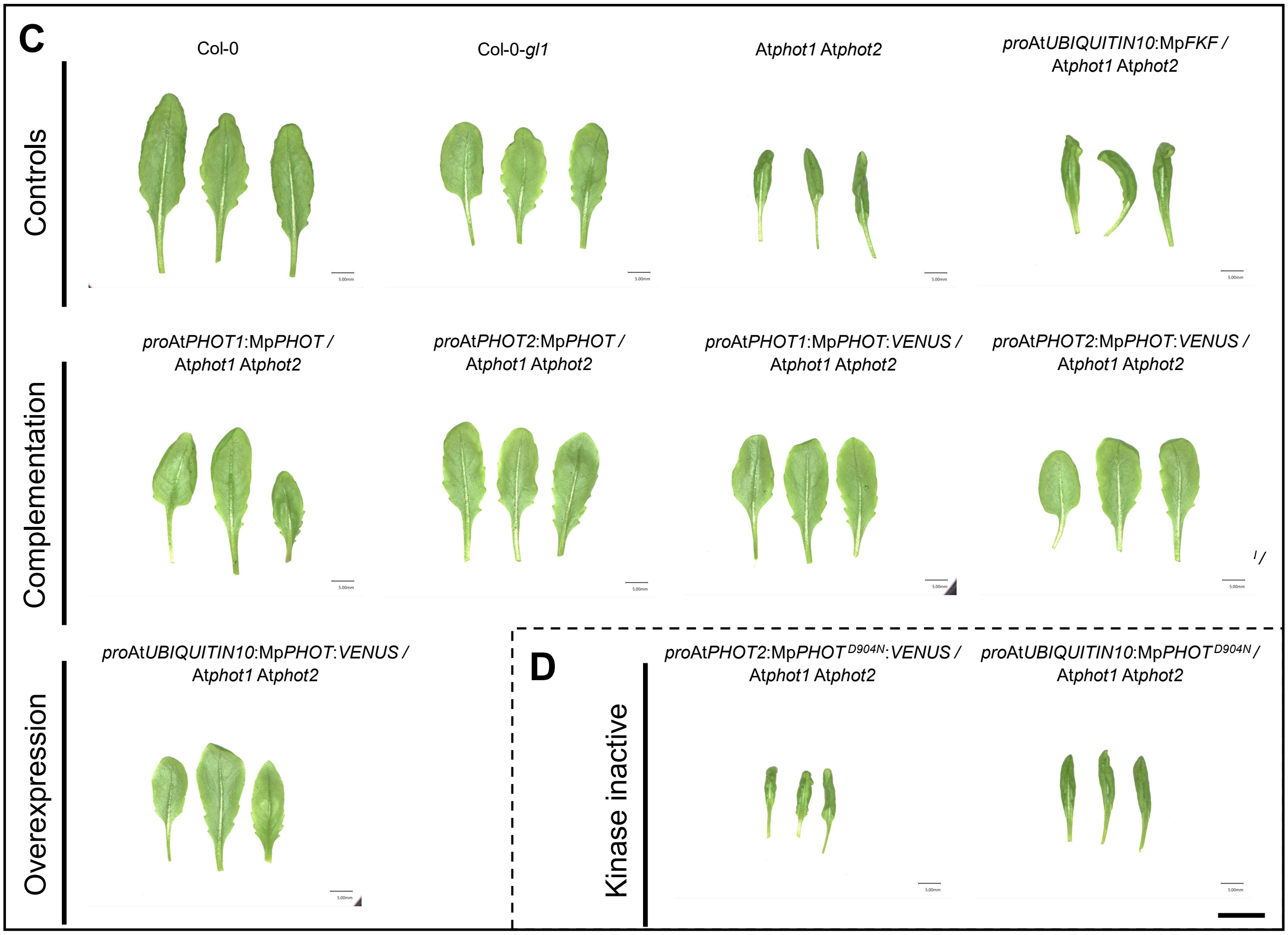
C
Controls
Col-0
Col-0-gl1
Atphot1 Atphot2
proAtUBIQUITIN10:MpFKF / Atphot1 Atphot2
Complementation
proAtPHOT1:MpPHOT / Atphot1 Atphot2
proAtPHOT2:MpPHOT / Atphot1 Atphot2
proAtPHOT1:MpPHOT:VENUS / Atphot1 Atphot2
proAtPHOT2:MpPHOT:VENUS / Atphot1 Atphot2
Overexpression
proAtUBIQUITIN10:MpPHOT:VENUS / Atphot1 Atphot2
D
Kinase inactive
proAtPHOT2:MpPHOT D904N:VENUS / Atphot1 Atphot2
proAtUBIQUITIN10:MpPHOT D904N / Atphot1 Atphot2

S3

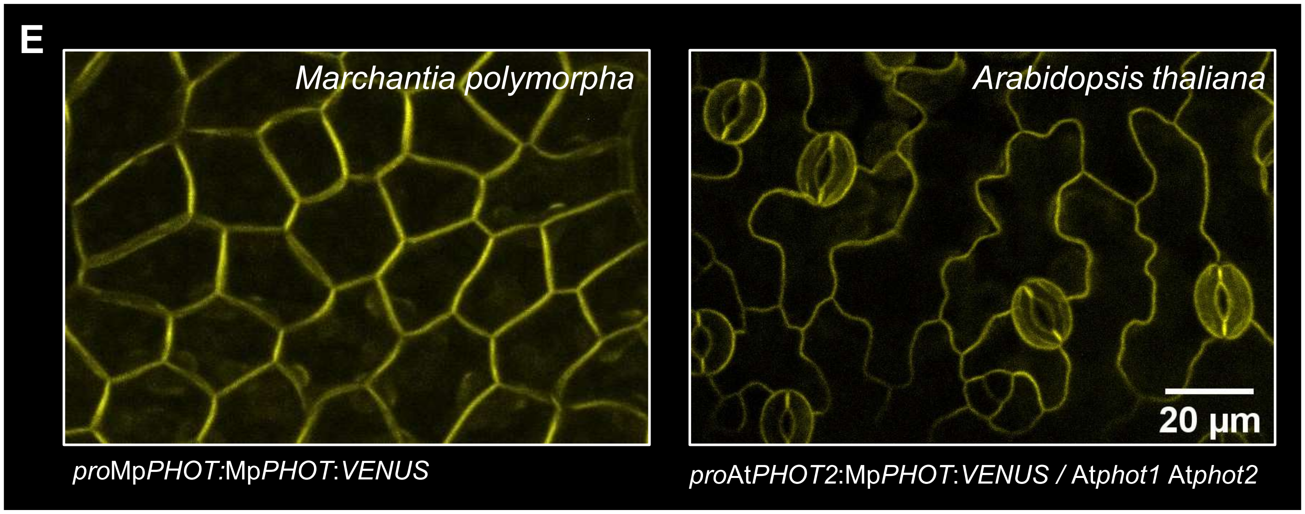
E
Marchantia polymorpha
Arabidopsis thaliana
20 µm
proMpPHOT:MpPHOT:VENUS
proAtPHOT2:MpPHOT:VENUS / Atphot1 Atphot2

S4

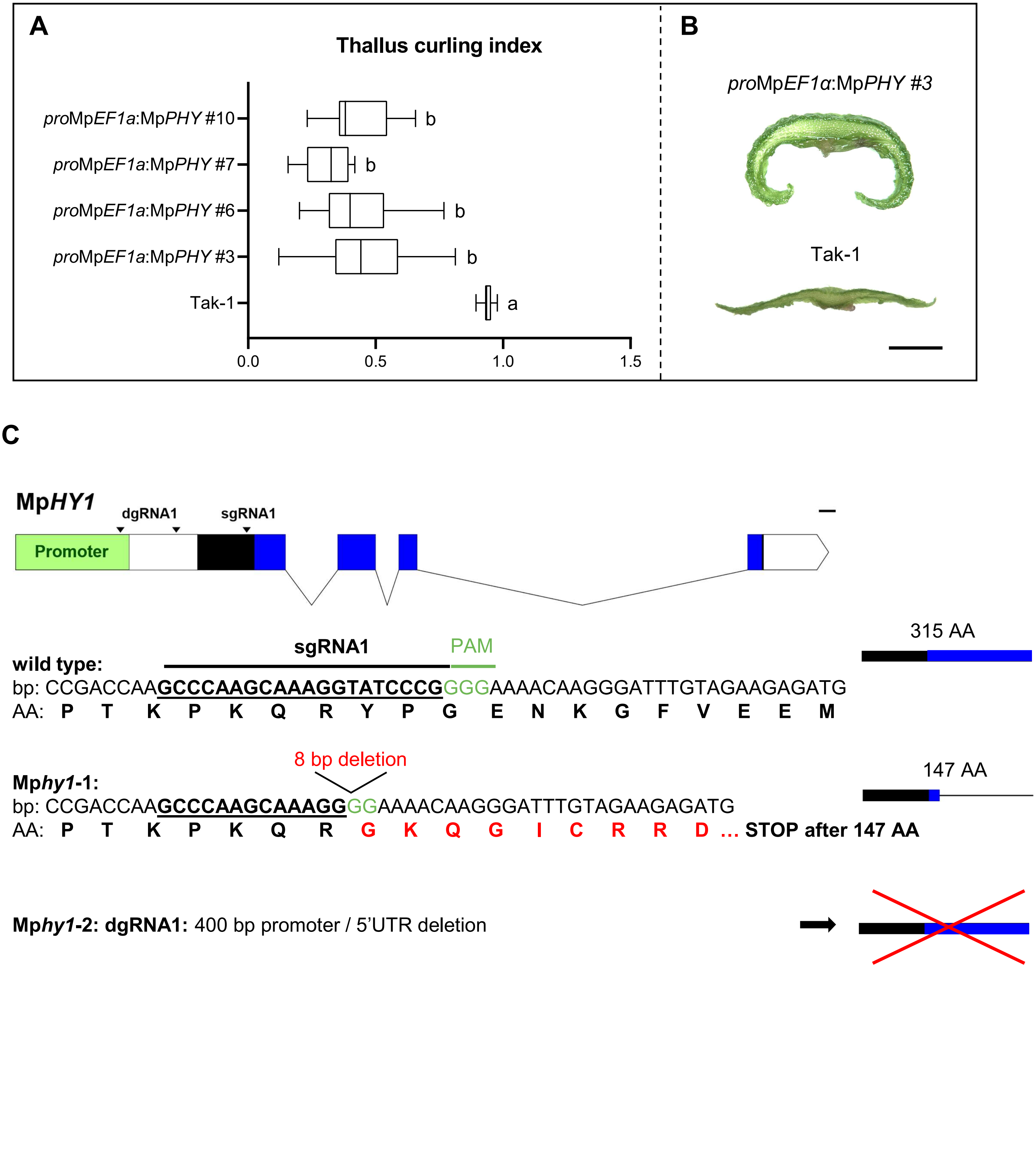
A
Thallus curling index
proMpEF1a:MpPHY #10
proMpEF1a:MpPHY #7
proMpEF1a:MpPHY #6
proMpEF1a:MpPHY #3
Tak-1
b
b
b
b
a
0.0
0.5
1.0
1.5
B
proMpEF1α:MpPHY #3
Tak-1
C
MpHY1
dgRNA1
sgRNA1
Promoter
315 AA
sgRNA1
PAM
wild type:
bp: CCGACCAAGCCCAAGCAAAGGTATCCCGGGGAAAACAAGGGATTTGTAGAAGAGATG
AA: P T K P K Q R Y P G E N K G F V E E M
8 bp deletion
147 AA
Mphy1-1:
bp: CCGACCAAGCCCAAGCAAAGGGGAAAACAAGGGATTTGTAGAAGAGATG
AA: P T K P K Q R G K Q G I C R R D … STOP after 147 AA
Mphy1-2: dgRNA1: 400 bp promoter / 5'UTR deletion

# S5

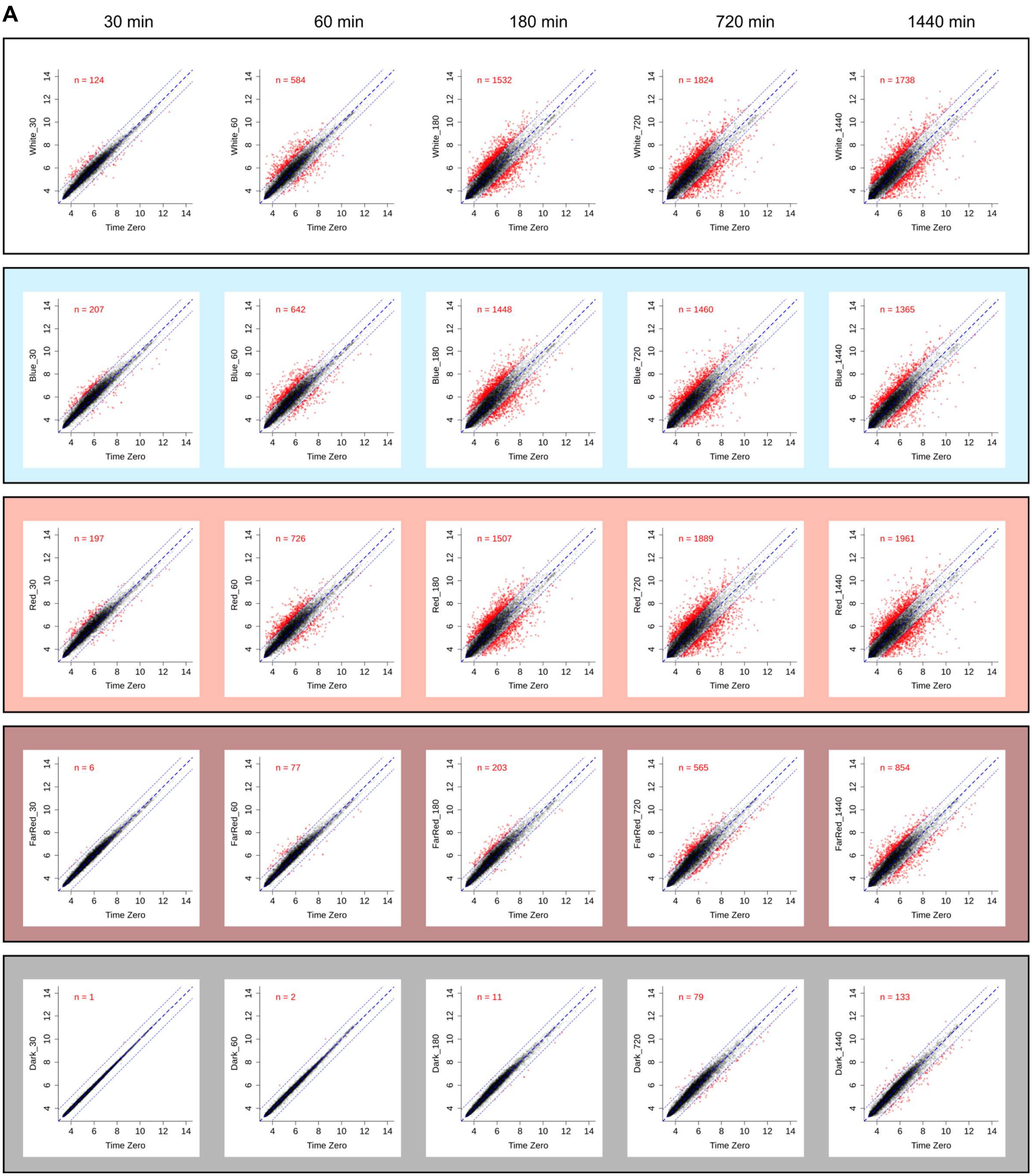
A
30 min
60 min
180 min
720 min
1440 min
n = 124
n = 584
n = 1532
n = 1824
n = 1738
White_30
White_60
White_180
White_720
White_1440
n = 207
n = 642
n = 1448
n = 1460
n = 1365
Blue_30
Blue_60
Blue_180
Blue_720
Blue_1440
n = 197
n = 726
n = 1507
n = 1889
n = 1961
Red_30
Red_60
Red_180
Red_720
Red_1440
n = 6
n = 77
n = 203
n = 565
n = 854
FarRed_30
FarRed_60
FarRed_180
FarRed_720
FarRed_1440
n = 1
n = 2
n = 11
n = 79
n = 133
Dark_30
Dark_60
Dark_180
Dark_720
Dark_1440
Time Zero

# S5

**B**

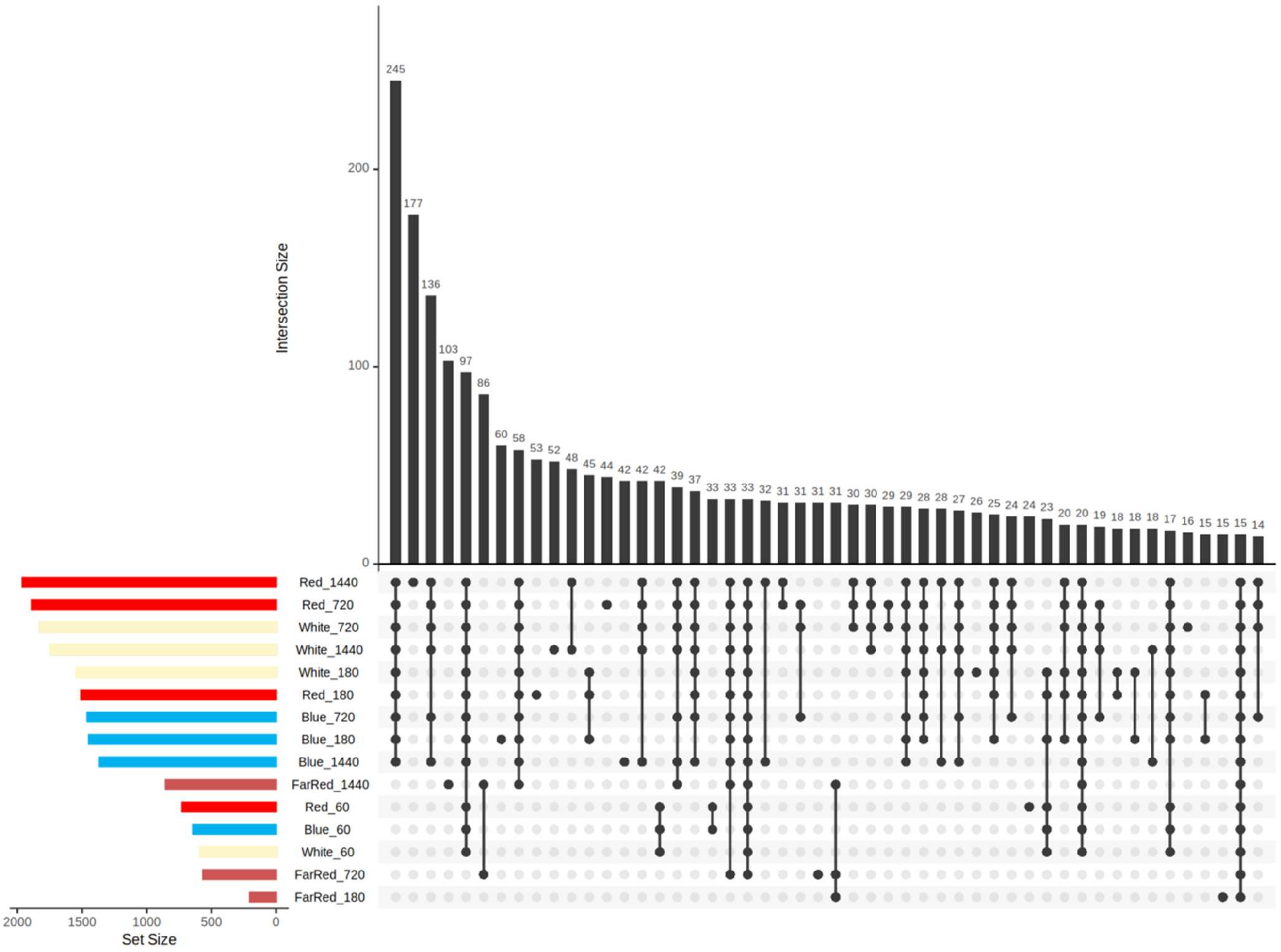

Intersection Size
245 177 136 103 97 86 60 58 53 52 48 45 44 42 42 42 39 37 33 33 33 32 31 31 31 31 30 30 29 29 28 28 27 26 25 24 24 23 20 20 19 18 18 18 17 16 15 15 15 14
0
100
200
Red_1440
Red_720
White_720
White_1440
White_180
Red_180
Blue_720
Blue_180
Blue_1440
FarRed_1440
Red_60
Blue_60
White_60
FarRed_720
FarRed_180
2000
1500
1000
500
0
Set Size


**C**

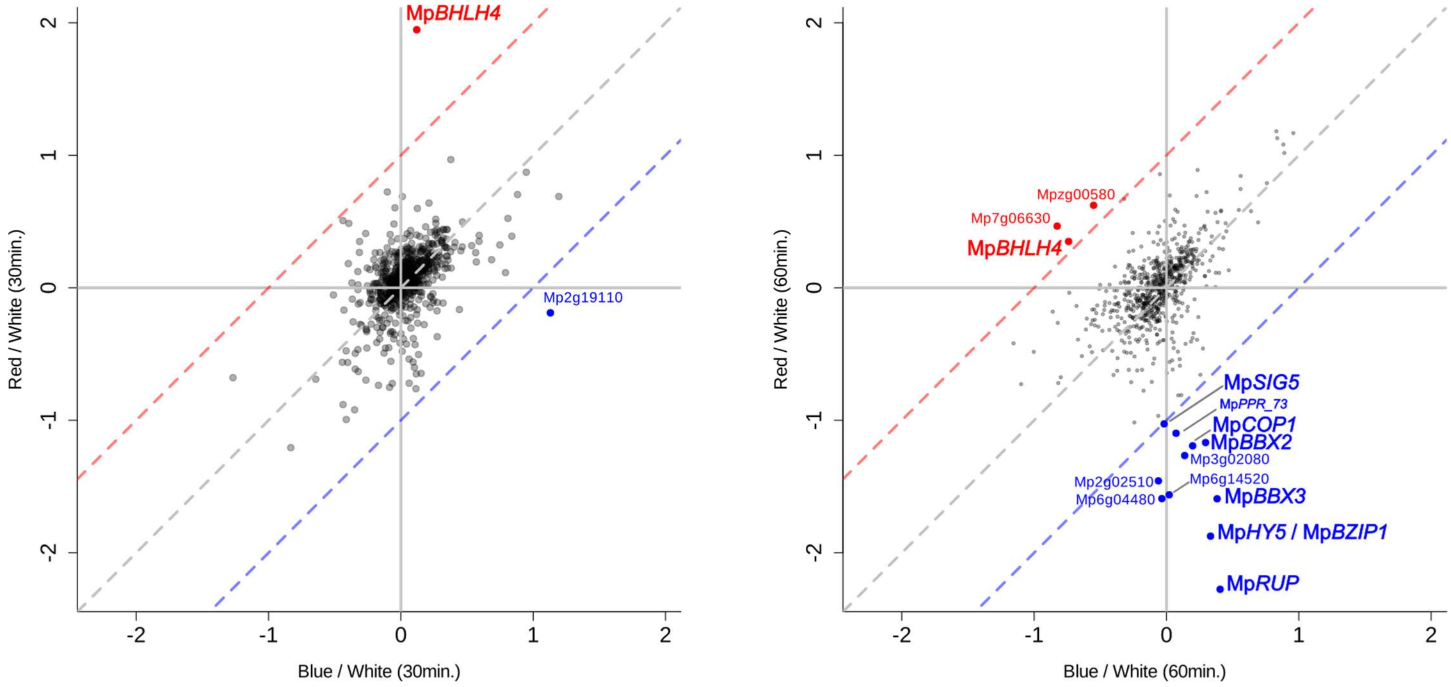

MpBHLH4
Mp2g19110
Red / White (30min.)
Blue / White (30min.)
-2
-1
0
1
2
Mpzg00580
Mp7g06630
MpBHLH4
MpSIG5
MpPPR_73
MpCOP1
MpBBX2
Mp3g02080
Mp2g02510
Mp6g14520
Mp6g04480
MpBBX3
MpHY5 / MpBZIP1
MpRUP
Red / White (60min.)
Blue / White (60min.)

**S6**

**A**

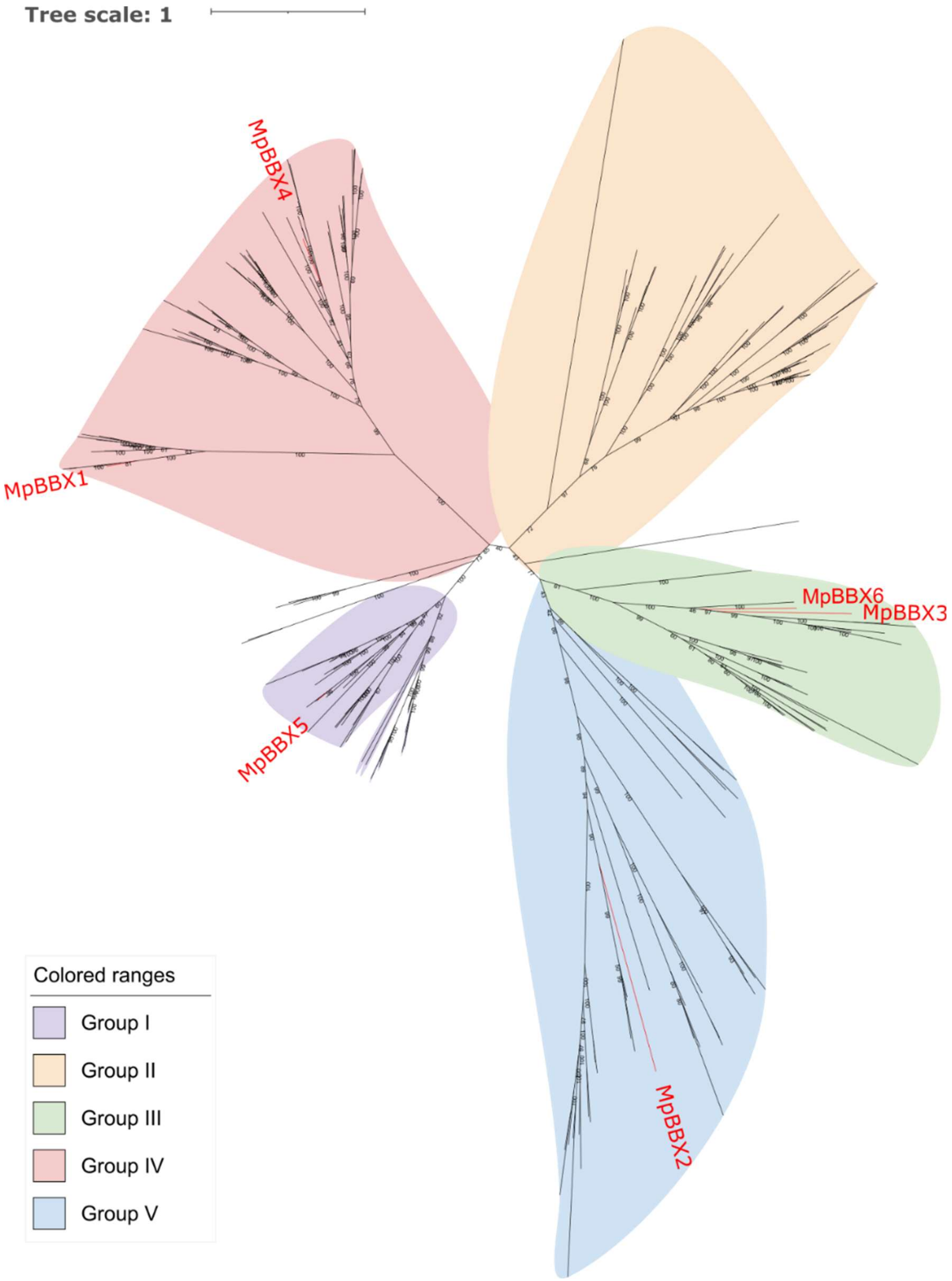
Tree scale: 1
MpBBX4
MpBBX1
MpBBX5
MpBBX6
MpBBX3
MpBBX2
Colored ranges
Group I
Group II
Group III
Group IV
Group V

# S6

**B**

**Mp*BBX1***

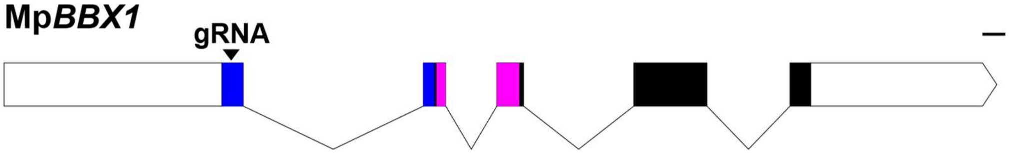


237 AA

**wild type:** gRNA PAM
bp: CCCGCGCG**ACTTTTTTGTGCCGCCGACG**AGGCTGCACTGTGCTTGACATGT
AA: **P A R L F C A A D E A A L C L T C**

23 AA

***bbx1*-1:** 6 bp and 17 bp insertion
bp: CCCGCGCG**ACTTTTTTGTGCCGCTGCAGGCAGGGTTTAGGGTTTAGGGACG**AGG
AA: **P A R L F C A A A G R V * STOP after 23 AA**

28 AA

***bbx1*-2:** 8 bp insertion
bp: CCCGCGCG**ACTTTTTTGTGCCGCCTTTTTTCTGACG**AGGCTGCACTGTGCTTGA
AA: **P A R L F C A A F F L T R L H C A * STOP after 28 AA**

21 AA

***bbx1*-3:** Large indel
bp: CCCGCGCG**ACTTTTTTGTGCCGCCGTCTTCTAGCAGTTCTTCTTCTAGAACTTCTTCTTTTTCACATGTCAAGCA**CTGCACTGTGCTTGACATGT
AA: **P A R L F C A A V F * STOP after 21 AA**

**Mp*BBX5***

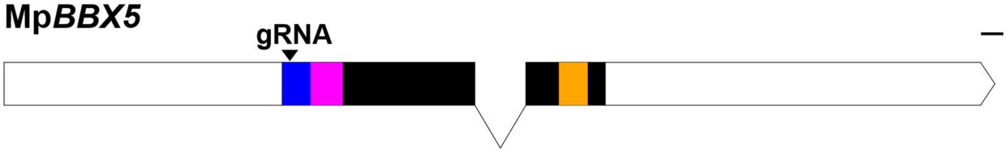


403 AA

**wild type:** gRNA PAM
bp: TCATC**GAGCGCAGTCGTCTACTGCC**GGGCCGACGCCGCCTTCCTGTGCGCAGGGTGTGACACC
AA: **S S S A V V Y C R A D A A F L C A G C D T**

28 AA

***bbx5*-1:** 1 bp insertion
bp: TCATC**GAGCGCAGTCGTCTACTTGCC**GGGCCGACGCCGCCTTCCTGTGCGCAGGGTGTGA
AA: **S S S A V V Y L P G R R R L P V R R V * STOP after 28 AA**

28 AA

***bbx5*-2:** 1 bp insertion
bp: TCATC**GAGCGCAGTCGTCTACTTGCC**GGGCCGACGCCGCCTTCCTGTGCGCAGGGTGTGA
AA: **S S S A V V Y L P G R R R L P V R R V * STOP after 28 AA**

55 AA

***bbx5*-3:** 7 bp deletion
bp: TCATC**GAGCGCAGTCGCC**GGGCCGACGCCGCCTTCCTGTGCGCAGGGTGTGACACC
AA: **S S S A V A G P T P P S C A Q G V T … STOP after 55 AA**

# S6

## B

##### Mp*BBX2*

gRNA PAM

**wild type:**

bp: CGTCTTTGTCC**CCGCTGCAACCCCTCCACGA**CGGCAGCAGTCGTGAACGAT

AA: **R L C P R C N P S T T A A V V N D**

***bbx2*-1:** 2 substitutions and 7bp insertion

bp: CGTCTTTGTTC**CCGCTGCAACCCCCCGACTGTTCACGA**CGGCAGCAGTCGTGAACGATTGCAACGGTGA

AA: **R L C S R C N P P T V H D G S S R E R L Q R * STOP after 89 AA**

***bbx2*-2:** 1 bp insertion

bp: CGTCTTTGTCC**CCGCTGCAACCCCTCCAACGA**CGGCAGCAGTCGTGAACGATTGCAACGGTGA

AA: **R L C P R C N P S N D G S S R E R L Q R * STOP after 87 AA**

##### Mp*BBX3*

gRNA 1 PAM gRNA 2 PAM

**wild type:**

bp: AAAACCGCA**ACAGCCGTGATGTCCATGGC**CGGACGGGCC**GTGCGTGCATGCGACGTCTG**CGGGCGCGACCGC

AA: **K T A T A V M S M A G R A V R A C D V C G R D R**

***bbx3*-1:** 1 bp insertion

bp: AAAACCGCA**ACAGCCGTGATGTCCATAGGC**CGGACGGGCCGTGCGTGCATGCGACGTCTGCGGGCGCGACCGC

AA: **K T A T A V M S I G R T G R A C M R R L R A R P…**

*** STOP after 196 AA**

***bbx3*-2:** 10 bp deletion

bp: AAAACCGCAACAGCCGTGATGTCCATGGCCGGACGGGCC**GTGCGTGCATGCGACGT**GACCGCG

AA: **K T A T A V M S M A G R A V R A C D V T A … * STOP after 67 AA**

##### Mp*BBX4*

gRNA PAM

**wild type:**

bp: ATGAGGGT**TCAGTGTGATGGATGTGAGA**GGGCCGTGGCGTCGATTATGTGTTGCGCGGATGAAGCT

AA: **M R V Q C D G C E R A V A S I M C C A D E A**

***bbx4*-1:**

bp: ATGAGGGT**TCAGTGTGATGGATGTGAAGA**GGGCCGTGGCGTCGATTATGTGTTGCGCGGATGA

AA: **M R V Q C D G C E E G R G V D Y V L R G * STOP after 20 AA**

***bbx4*-2:**

44bp deletion (deleting part of 5’UTR, start codon and gRNA target site)

bp: (ATGAGGGT**TCAGTGTGATGGATGTGAGA**GGGCCGTGGC)GTCGATTATGTGTTGCGCGGATGAAGCT

AA: **M R V Q C D G C E R A V A S I M C C A D E A**

##### Mp*BBX6*

gRNA PAM

**wild type:**

bp: GCCGACGAAGCG**TATCTGTGCGAGCCGTGCGA**TGGGTCCGTGCACAACGCGAATGCT

AA: **A D E A Y L C E P C D G S V H N A N A**

***bbx6*-1:** 1 bp insertion, 1bp substitution

bp: GCCGACGAAGCG**TATCTGTGCGAGCCGTGACGA**TGGATCCGTGCAC

AA: **A D E A Y L C E P * STOP after 47 AA**

***bbx6*-2:** 13 bp deletion

bp: GCCGACGAAGCG**TATCTGTGCGAGCCGTGC**ACAACGCGAATGCT

AA: **A D E A Y L C E P C T T R M … * STOP after 69 AA**

S6

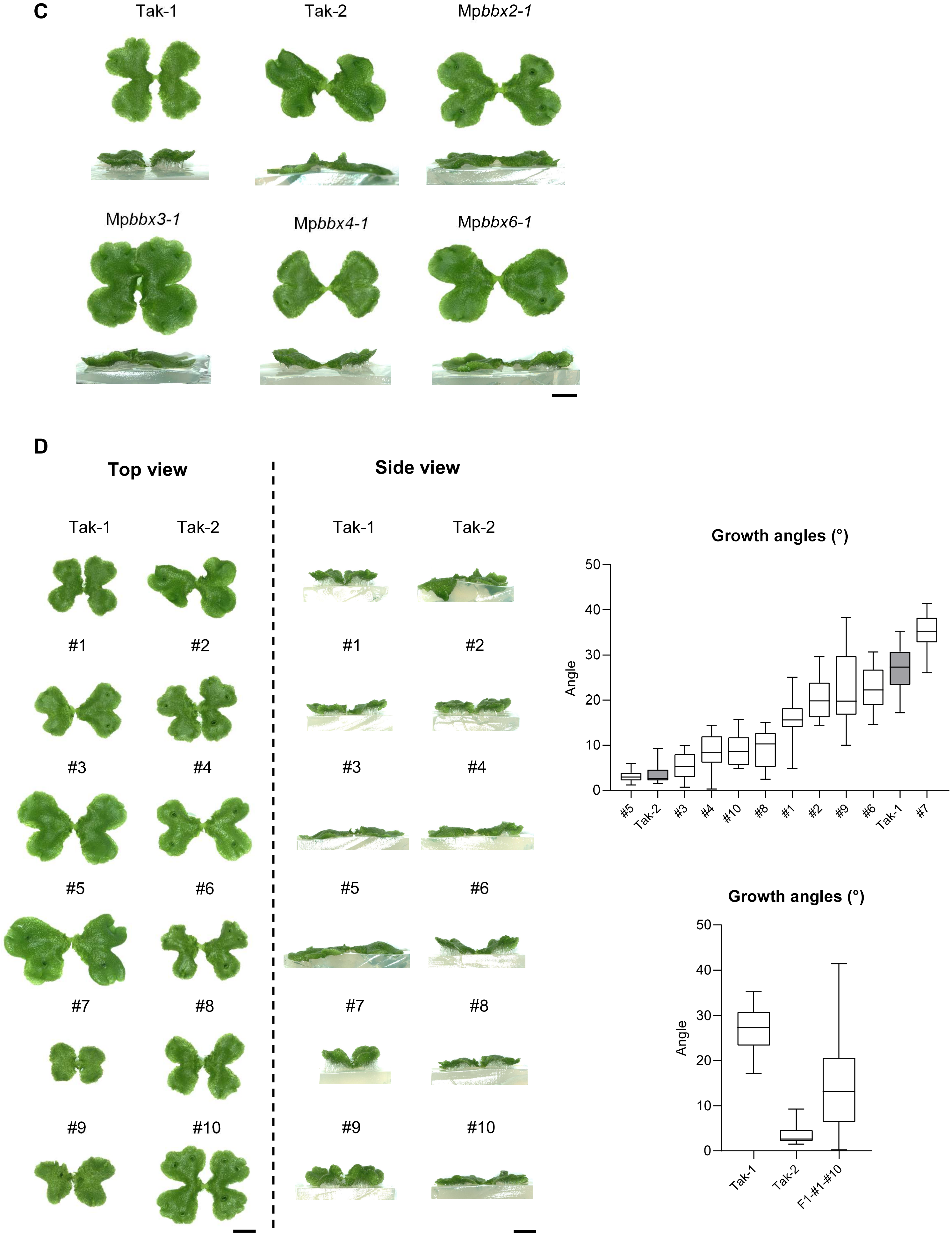
C
Tak-1
Tak-2
Mpbbx2-1
Mpbbx3-1
Mpbbx4-1
Mpbbx6-1
D
Top view
Side view
Tak-1
Tak-2
Tak-1
Tak-2
#1
#2
#3
#4
#5
#6
#7
#8
#9
#10
Growth angles (°)
Angle
50
40
30
20
10
0
#5
Tak-2
#3
#4
#10
#8
#1
#2
#9
#6
Tak-1
#7
Growth angles (°)
Angle
50
40
30
20
10
0
Tak-1
Tak-2
F1-#1-#10
